# Diverging Neural Dynamics for Syntactic Structure Building in Naturalistic Speaking and Listening

**DOI:** 10.1101/2022.10.04.509899

**Authors:** Laura Giglio, Markus Ostarek, Daniel Sharoh, Peter Hagoort

## Abstract

The neural correlates of sentence production have been mostly studied with constraining task paradigms that introduce artificial task effects. In this study, we aimed to gain a better understanding of syntactic processing in spontaneous production vs. naturalistic comprehension. We extracted word-by-word metrics of phrase-structure building with top-down and bottom-up parsers that make different hypotheses about the timing of structure building. In comprehension, structure building proceeded in an integratory fashion and led to an increase in activity in posterior temporal and inferior frontal areas. In production, structure building was anticipatory and predicted an increase in activity in the inferior frontal gyrus. Newly developed production-specific parsers highlighted the anticipatory and incremental nature of structure building in production, which was confirmed by a converging analysis of the pausing patterns in speech. Overall, the results showed that the unfolding of syntactic processing diverges between speaking and listening.

## Introduction

Studies on the neurobiology of language typically use highly controlled experimental paradigms far removed from typical language experience in everyday life. However, in the last decade, there has been a shift towards naturalistic studies of language processing. This shift has happened in multiple directions, from the use of virtual environments to increase ecological validity (Huizeling et al., 2021; Peeters, 2019), to the auditory presentation of audiobooks or narrative reading with neuroimaging (Alday et al., 2017; Brennan, 2016; Heilbron et al., 2022; Willems & Gerven, 2018). The increased ecological validity in naturalistic studies opens a window into language processing free of the artificiality of task designs, whose main goal is to isolate specific features of language (Andric & Small, 2015). In traditional settings, experimental control comes at the cost of context, which is heavily reduced to minimize confounds. This contrasts with the highly contextual nature of everyday language use, creating a large gap between the actual object of study and its realization in experiments. Combining naturalistic stimuli and advanced analysis methods, such as audiobooks and probabilistic parsers, has the potential to bring the two much closer together (Brennan, 2016; Hale et al., 2022).

The overwhelming majority of studies on the neurobiology of language is on comprehension, while speaking is largely unexplored. Importantly, while naturalistic studies are becoming more common in the field of language comprehension, studies of naturalistic *production* are still lacking. This is problematic because the gulf between spontaneous production and production in controlled experiments is particularly large. In spontaneous language production, the speaker is by definition in control of what to say. In contrast, experimental paradigms attempt to have as much control over participants’ speech as possible. This has usually been achieved with picture description experiments or with the use of visual probes together with written linguistic stimuli (e.g. Giglio et al., 2022; Griffin & Bock, 2000; Matchin & Hickok, 2016; Takashima et al., 2020). While these strategies have allowed for controlled investigations of linguistic processing, they may be confounded with the heavy task requirements that make controlled production very removed from everyday speaking.

In this fMRI study, we aimed to study syntactic processing in spontaneous production and comprehension in order to understand whether and how they differ. Previous studies found shared neural resources for production and comprehension (Menenti et al., 2011; Segaert et al., 2012) and a similar network for processing syntactic complexity across modalities (Giglio et al., 2022; Hu et al., 2022). At the same time, production was found to elicit larger responses to syntactic complexity than comprehension, especially in the left inferior frontal gyrus (LIFG) (Giglio et al., 2022; Hu et al., 2022; Indefrey et al., 2004). The differential sensitivity to complexity between modalities may be due to two main factors: 1) Speaking is a form of action, unlike the more passive process of listening. The message needs to be fully and correctly encoded into a linear sequence that results in articulation (Bock, 1982; Garrett, 1980; Garrett, 1982, but see Goldberg & Ferreira, 2022). In comprehension, instead, one may choose how much attention to pay to the linguistic input, and sometimes may fail to correctly parse ambiguous input (Ferreira, 2003; Ferreira et al., 2002; for discussion, see Ferreira & Lowder, 2016). 2) Alternatively, previous studies may have been affected by unequal task requirements between modalities. The tasks used to elicit controlled production are usually more artificial than the respective tasks in comprehension, where only listening is expected, even if the same stimuli are used. To reduce task differences, here we focused on the neural response of syntactic processing in spontaneous production and compared it to the neural response to comprehension of the same materials. To the best of our knowledge, this is the first fMRI study of syntactic processing with unconstrained speaking.

To model linguistic processing in spontaneous production, we used word-by-word indices that were previously used to successfully predict brain activity in comprehension (e.g. Brennan et al., 2016; Lopopolo et al., 2021; Nelson et al., 2017). Widely used continuous indexes of linguistic processing include measures of word surprisal that show the brain’s sensitivity to predictability (Henderson et al., 2016; Shain et al., 2020; Willems et al., 2016); continuous measures of syntactic tree building (Brennan et al., 2016, 2020; Brennan & Hale, 2019; Lopopolo et al., 2021; Nelson et al., 2017; Stanojević et al., 2021); and word embeddings modelling semantic features (Wehbe et al., 2014). Focusing on syntactic processing, increasingly sophisticated approaches that characterize continuous structure building have highlighted that left fronto-temporal regions are sensitive to measures of syntactic processing (Bhattasali et al., 2019; Brennan et al., 2016, 2020; Li & Hale, 2019; Nelson et al., 2017; Stanojević et al., 2021). We asked how syntactic processing, modelled with continuous metrics of syntactic tree building, affects brain activity in production and comprehension.

In the current study, we compared two parser models, a top-down and a bottom-up parser strategy. These strategies account for the same structure but make different hypotheses about the timing of syntactic operations. Top-down parsers build nodes at phrase-opening in an anticipatory fashion, whereas bottom-up parsers build nodes at phrase-closing in an integratory fashion. Here, we hypothesized the timing of operations to be the critical difference between production and comprehension, due to the different requirements and inputs of each modality (Momma & Phillips, 2018). In production, the speaker has some knowledge about the upcoming structure, since the structure related to the words that are uttered must have been computed (Bock & Levelt, 1994; Levelt, 1989). In comprehension, instead, after accounting for predictable continuations, listeners need to wait for the input to fully compute the structure. We thus expected the more anticipatory top-down parser to better predict neural activity in production and bottom-up parsing to better predict neural activity in comprehension. In a follow-up exploratory analysis, we explored whether alternative parsing strategies may be more fitting for production, since the parser models discussed so far were developed for language processing in comprehension specifically. In particular, we developed two parsers that assume different levels of incremental processing, by making different predictions about how early phrase-structure building operations occur.

Finally, we investigated which regions responded to syntactic processing load in each modality. In particular, we focused on two parts of the LIFG (*pars opercularis* or BA44, and *pars triangularis* or BA45) and on the left posterior middle temporal gyrus (LpMTG). These regions were all found to be responsive to syntactic manipulations in both modalities (Hagoort & Indefrey, 2014; Lopopolo et al., 2021; Pallier et al., 2011; Snijders et al., 2009; Takashima et al., 2020; Zaccarella, Meyer, et al., 2017; Zaccarella, Schell, et al., 2017), sometimes with differences in their sensitivity to each modality (Giglio et al., 2022; Indefrey, 2018; Matchin & Wood, 2020). In particular, the LIFG was found to be more responsive to syntactic complexity in production than comprehension (Giglio et al., 2022; Indefrey et al., 2004), while the LpMTG was more responsive during comprehension (Giglio et al., 2022; Matchin & Wood, 2020). We thus investigated to what extent these regions are sensitive to continuous metrics of syntactic processing, and whether the modality differences observed in earlier studies (Giglio et al., 2022; Indefrey, 2018; Matchin & Wood, 2020) are replicated in naturalistic settings.

## Results

### Incremental metrics of phrase-structure building

To obtain incremental metrics of syntactic processing, we proceeded in two steps. First, we extracted the constituent structure of each sentence with a probabilistic context-free phrase-structure grammar (Stanford parser, Klein & Manning, 2003). From the extracted constituent parse, we then computed the parser operations carried out at each word according to different parsing models (Hale, 2014). These parsers incrementally build the syntactic structure of a sentence following different strategies, leading to a hypothesized number of phrase-structure building operations that need to be carried out at each word (Hale, 2014). This results in an incremental complexity metric that corresponds to the number of nodes that are built with each word. A top-down strategy builds the phrase structure from the top of the tree to a given word, such that it predicts increased activity when phrases are opened. In comprehension, it sometimes anticipates nodes before they are unambiguous to the listener, for example in the presence of adjuncts. Bottom-up parsing instead builds the phrase structure only after all the evidence has been seen, that is, after all words attached to each node have been met. It thus predicts increased activity when phrases are closed. Ultimately, both strategies lead to the same node count, but they make different predictions about the timing of syntactic operations and thus of corresponding neural activation (see Methods for more details, Fig.1A-B).

**Figure 1:**
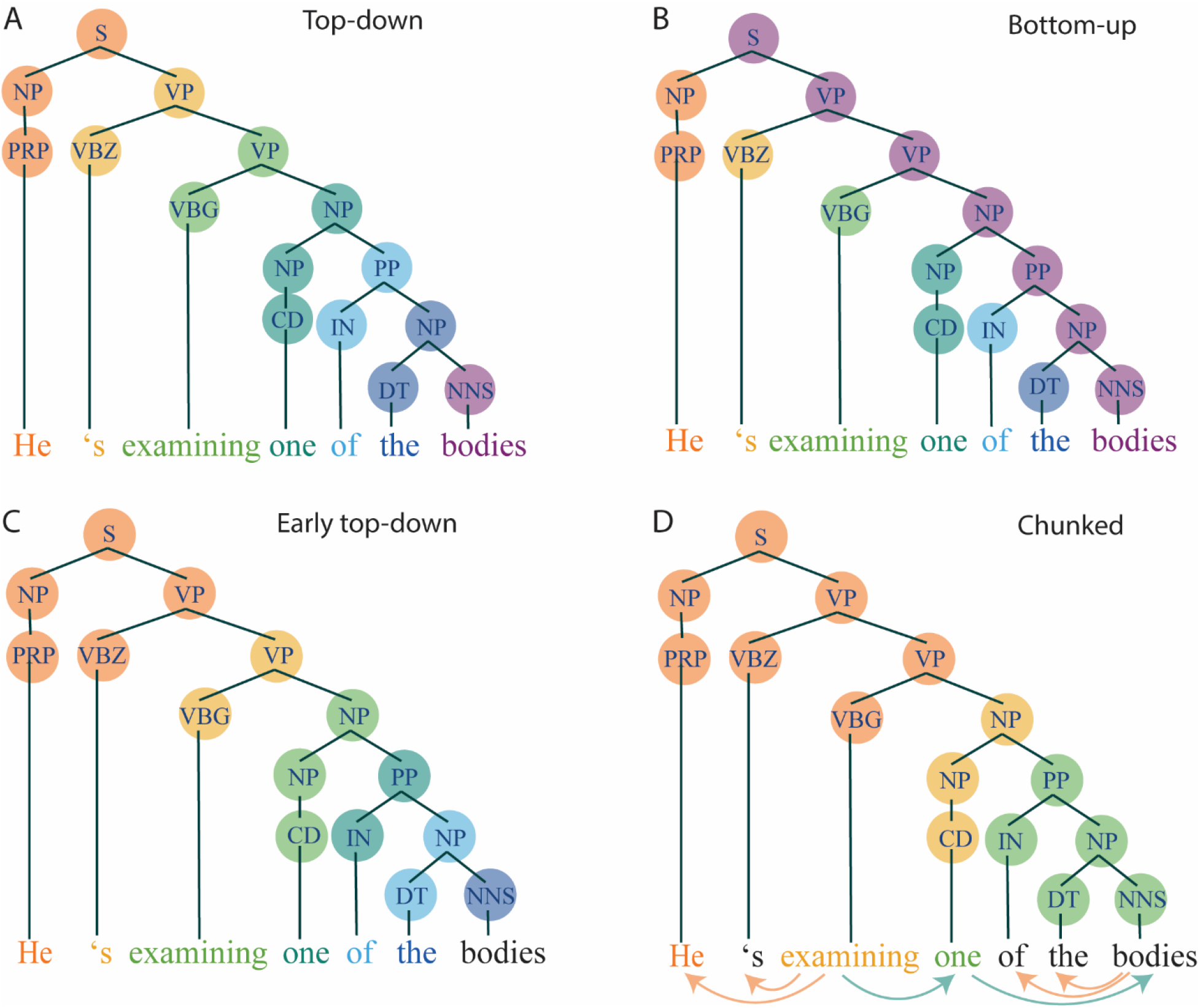
Node counting following different parsing strategies. The colored circles refer to the nodes that are built at the time point the word in the same color is uttered or heard. **A**: Colored representation of top-down phrase structure building, with nodes counted from the top of the tree to the word. **B**: Colored representation of bottom-up phrase structure building, with nodes counted from the bottom of the tree (i.e. the terminal nodes) to the top. Only nodes where both daughter nodes have been already met can be counted at each word. **C**: Colored representation of early top-down phrase-structure building, assuming operations to take place before word onset (production-specific). **D**: Colored representation of chunked phrase-structure building, following a less incremental strategy (production-specific). This node counting strategy is chunked based on the heads of the dependency parse of the same sentence (shown by the arrows below words, also see Supplementary Fig.1). Heads are words from which an arrow originates. The nodes of the same constituent structure used by the other strategies are counted here. The chunked nature of this parser results in phrase-structure building operations to be assigned to some but not all words in a sentence. Black words are words that are not assigned any phrase-structure building operation (e.g. sentence-final words).

We also quantified the load of processing complexity on working memory with an *open nodes* measure. This measure counts the number of nodes that have been opened (i.e. counted by the top-down strategy) but have not been closed yet (i.e. counted by the bottom-up strategy), tracking the numbers of words that need to be kept in working memory until they can be merged in a constituent (Nelson et al., 2017). In other words, this complexity metric tracks how much of the hypothesized structure needs to be confirmed by upcoming input. We expected this index to predict an increase in activity in comprehension, following Nelson et al. (2017) and Uddén et al. (2019). In production, it would also lead to an activity increase if speakers kept track of the structure that needs to be closed. Finally, to make sure that the syntactic predictors did not simply track word probabilities based on context, we quantified word surprisal from transformer model GPT-2 (Radford et al., 2019).

### Distinct dynamics for phrase-structure building in language production vs. comprehension

To directly compare the word-by-word predictors with BOLD activity with a 1.5 second resolution (thus including several words at each fMRI volume), we convolved the linguistic predictors with the haemodynamic response function and resampled it to the 1.5 repetition time (see Methods for more details, Fig. 2). We then regressed average BOLD activity in BA44, BA45 and LpMTG in subject space against the predicted timeseries for each linguistic predictor with linear mixed-effects models.

**Figure 2:**
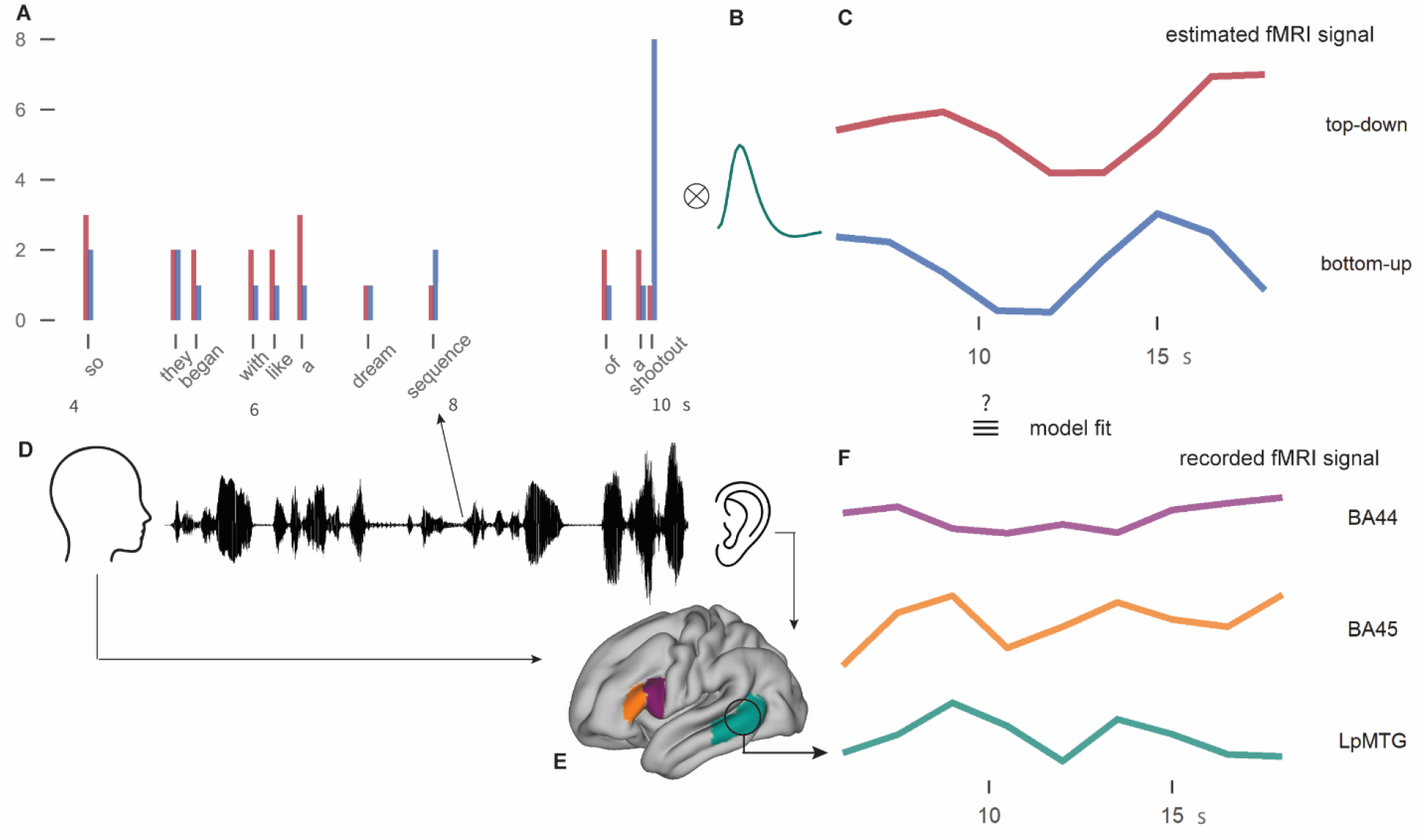
Graphical representation of the analysis paradigm to relate word-by-word predictors of linguistic complexity to BOLD activity. **A**: Word-by-word predictors of syntactic complexity were extracted from the constituent structure of the sentence spoken by a participant and listened to by other participants (**D**). The height of the bars in **A** represents the number of phrase-structure building operations expected to take place at each word following top-down and bottom-up parsing strategies (e.g. at “so” 3 nodes are counted for top-down, 2 for bottom-up). The weights of the syntactic predictors were convolved with the haemodynamic response function (**B**) to get predictor timeseries of BOLD activity at 1.5 sec resolution (**C**). These predictors timeseries were then compared to the average BOLD activity (**F**) in the brain of the speaker or the listener (**D**) in the three regions of interest (BA44, BA45 and LpMTG, **E**).

First, we ran a linear model of phrase-structure building operations on neural activity in BA44, BA45 and LpMTG (Fig. 3). The model included word rate, syllable rate, word frequency, word surprisal, top-down, bottom-up and open nodes, language modality and ROI (Supplementary Table 1). Word rate, word frequency and word surprisal significantly predicted an increase in BOLD activity (βs > 0.11, *t*s > 2.5, *p*s < 0.015). Syllable rate also marginally significantly predicted an increase in BOLD activity. More words, longer and less predictable words thus led to an increase in BOLD, while less frequent words, which should be harder to process, led to a decrease in activity. The effect of modality was also marginally significant (β = 0.008, SE = 0.004, *t* = 1.8, χ^2^ = 3.3, *p* < 0.068), with production having more positive activity than comprehension. The effect of modality did not interact with the effect of ROI.

**Figure 3:**
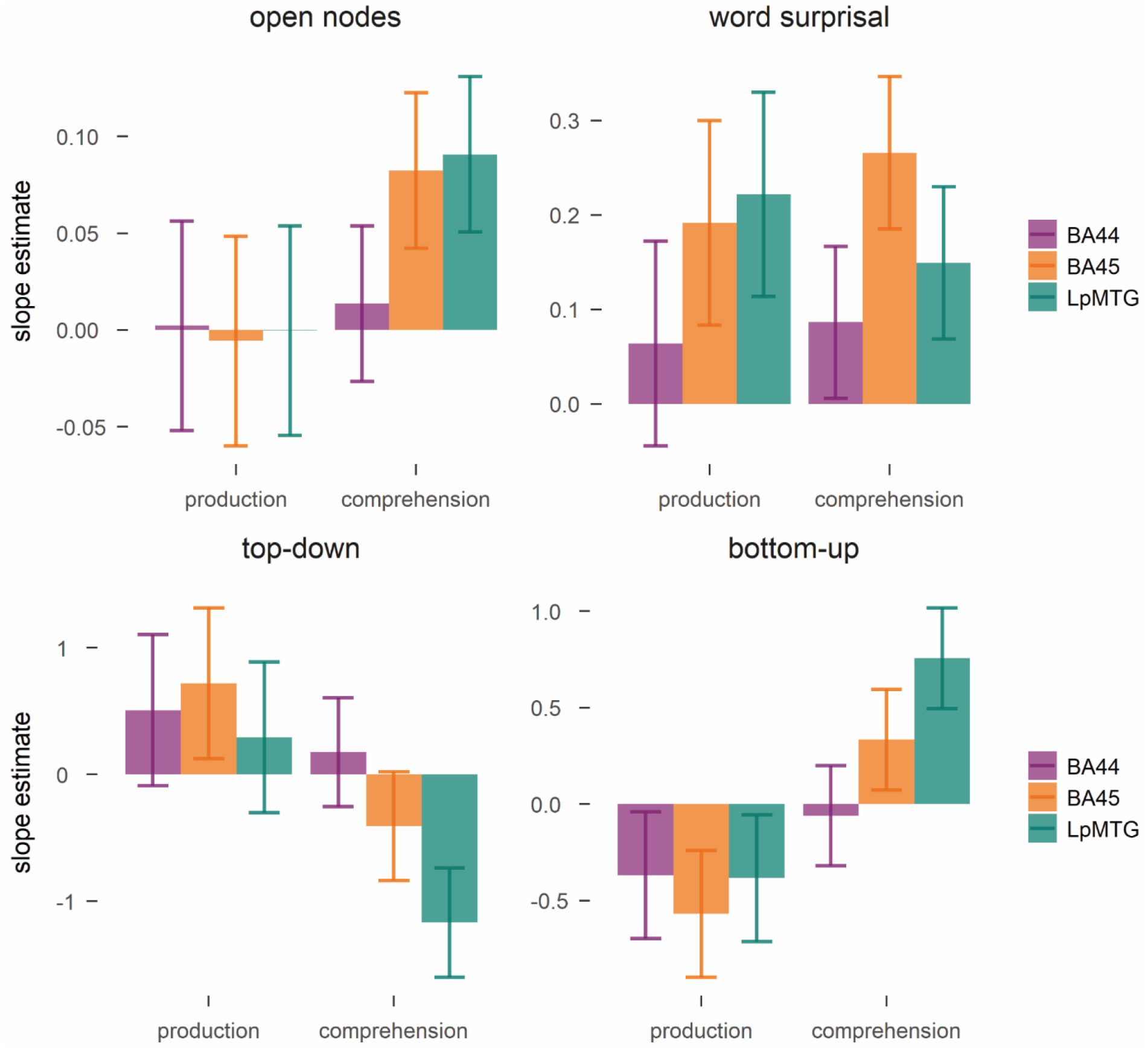
Beta estimates for the effect of each predictor of phrase-structure building, open nodes and word surprisal on BOLD activity in the regions of interest. Error bars represent 95% confidence intervals.

Larger word surprisal elicited an increase in BOLD in both modalities (Fig. 3, β = 0.16, SE = 0.02, *t* = 6.6, χ^2^ = 50.7, *p* < 0.0001). This effect interacted with ROI (β = 0.08, SE = 0.02, *t* = 3.6, χ^2^ = 15.4, *p* < 0.0005), since BA44 responded significantly less to surprisal than BA45 and LpMTG (estimates > 0.1, *p* < 0.03) in both modalities. Open nodes also had a significant effect on BOLD activity (Fig. 4, β = 0.031, SE = 0.015, *t* = 2.1, χ^2^ = 8.9, *p* < 0.003). The effect interacted with modality and ROI (χ^2^ = 8.9, *p* < 0.015). It was significant only in comprehension in BA45 and LpMTG (estimates > 0.84, *p* < 0.001), while the estimates approached zero in all ROIs in production. Open nodes track the number of nodes to be kept in working memory until they can be integrated. It thus seems that the amount of structure that needs to be kept in working memory to be confirmed with the input leads to a brain activity increase in comprehension, but not in production.

**Figure 4:**
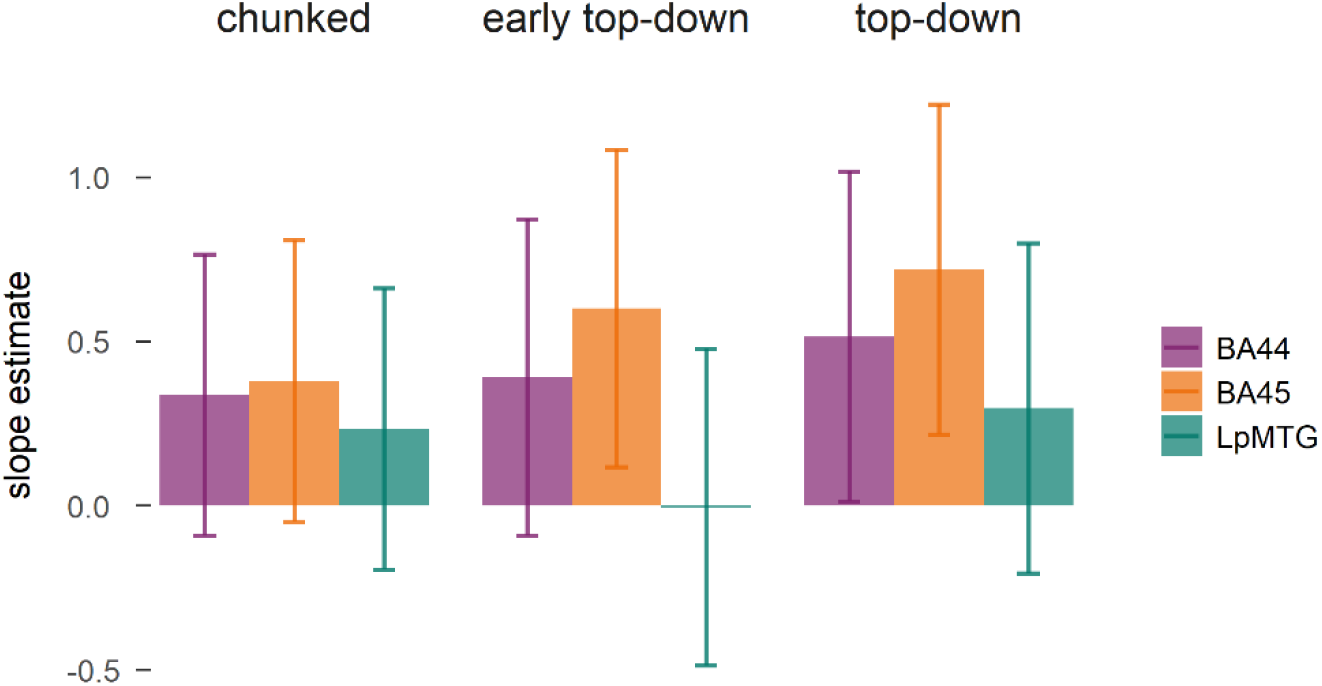
Beta estimates of the effect of each predictor of phrase-structure building in production on BOLD activity in the regions of interest. Error bars represent confidence intervals. The early top-down model led to the best model fit (Akaike information criterion, lower values indicate better fit): early top-down, 170803.9; top-down, 170821.3; chunked, 170834.5.

We next determined whether incremental metrics of phrase-structure building significantly predicted brain-activity in BA44, BA45 and LpMTG. Both top-down and bottom-up parsers added significant contributions to the model, in interaction with modality and ROI (two-way interactions: top-down, χ^2^ = 10.4, *p* < 0.006, bottom-up, χ^2^ = 12.7 *p* < 0.002). Anticipatory top-down node counts predicted a significant increase in activity in production only in BA45 (estimate = 0.49, SE = 0.2, *p* < 0.02), while they predicted a decrease in comprehension in BA45 and LpMTG (BA45: estimate = 0.28, SE = 0.15, *p* = 0.063, LpMTG: estimate = 0.8, SE = 0.15, *p* < 0.0001). The response to top-down node counts was significantly different among all ROIs in comprehension (difference estimates > 0.58, SE= 0.23, *p*s < 0.03), but not in production (difference estimates < 0.5, SE= 0.28, *p*s > 0.27). Integratory bottom-up node counts predicted an opposite pattern of results. Larger bottom-up counts led to a significant decrease in activity in production in all ROIs (estimates > 0.44, SE = 0.2, *p* < 0.03), while they predicted an increase in comprehension in BA45 and LpMTG (BA45: estimate = 0.4, SE = 0.16, *p* = 0.012, LpMTG: estimate = 0.9, SE = 0.15, *p* < 0.0001). Again, ROIs responded differently to bottom-up counts across modalities. In comprehension, the strongest response was in LpMTG (difference estimates > 0.39, SE= 0.16, *p*s < 0.035), while in production the responses were not significantly different among ROIs (estimates < 0.21, SE= 0.18, *p*s > 0.5). Therefore, activity in BA45 increased for different dynamics of phrase-structure building in production and comprehension, while the LpMTG positively responded only to comprehension. Activity in LpMTG and BA44 instead did not respond to anticipatory structure building in production, but decreased for integratory structure building, suggesting that both BA44 and LpMTG were active during sentence production, although not at the latencies predicted by the top-down parser, but their activity decreased at phrase-closing.

The parsers thus revealed marked differences between language production and comprehension. Anticipatory node counts led to an increase during production, but a decrease during comprehension, suggesting that in production syntactic structure building dominates at phrase opening. In comprehension, instead, neural activity may be reduced at phrase opening and may instead increase after phrase opening, when the top-down parser predicts a decrease in activity. This interpretation is confirmed by the activity increase in comprehension with integratory node counts, that predict increased activity at phrase closing. The decrease in activity for bottom-up counts in production suggests that at phrase closing syntactic processing load is reduced. Neural activity thus increases with syntactic structure building in both modalities, although with different dynamics, in BA45 in both production and comprehension, in LpMTG in comprehension only.

### Phrase-structure building in production proceeds in a highly incremental fashion

The parsing strategies mentioned so far were developed for comprehension. This is problematic because these parsers assign linguistic operations *at the time* a word is said. This is a reasonable assumption in comprehension, where processing follows the input. However, in production, once the word is articulated, the associated grammatical and lexical encoding has to have taken place already (e.g. Bock & Levelt, 1994; Indefrey & Levelt, 2004). We thus explored two production node building strategies that could better account for the timing of syntactic encoding in production. In both, the syntactic structure related to a word was assumed to be built at the latest while the previous word was articulated, but the two strategies made different predictions about the incrementality of structure building. An *early top-down* model predicted structure building to happen as the previous word is uttered (i.e. at each word we counted the nodes associated with the following word, see Methods for more details, Fig. 1C). This strategy leads to an equally incremental node building strategy as the original top-down strategy, but that, critically, happens earlier, more in line with theories of word production (Indefrey, 2011; Indefrey & Levelt, 2004).

While more fitting for production in terms of timing, this view presupposes a highly incremental syntactic encoder. However, studies of sentence planning in production have long debated whether planning is linearly or hierarchically incremental, that is, whether the structure is built from each concept separately or from the relations between concepts (Bock & Ferreira, 2014). Hierarchical models of sentence production consider the verb to be the central node for the syntactic structure (Bock & Levelt, 1994; Levelt, 1989), suggesting that planning proceeds less incrementally. We thus explored whether a less incremental parser would better account for brain activity than a word-by-word incremental parser. We developed a node building strategy that counted all the nodes between words that were identified as heads according to dependency parsing (see Methods for more details, Fig.1D and Supplementary Fig. 1). This *chunked* strategy predicts chunks of syntactic processing to happen at focal points, in a less incremental way.

We compared the initial top-down parser used in the previous analyses with the *early top down model* and the *chunked model* by fitting three linear mixed models to the production data, each with of these different predictors of phrase-structure building. The *early top-down* model led to the best model fit (as measured with the Akaike information criterion, lower values indicate better fit: *early top-down*, 170803.9; *top-down*, 170821.3; *chunked*, 170834.5, Supplementary Tables 2-4).

Interestingly, *top-down* and *chunked* models predicted an overall increase in BOLD (*top-down*, χ^2^ = 7.5, *p* < 0.007; *chunked*, χ^2^ = 3.9, *p* = 0.049), while the *early top-down* main effect was not significant (χ^2^ = 2.7, *p* = 0.1), but it interacted with ROI (χ^2^ = 6.2, *p* < 0.05) (Fig. 4). In particular, *early top-down* counts predicted an increase in BA45 (estimate = 0.44, SE = 0.18, *p* = 0.015), while the effect was absent in LpMTG (estimate = 0.003, SE = 0.18, *p* > 0.98). The estimate for LpMTG thus decreased when phrase-structure building operations were posited to take place earlier, suggesting that the LpMTG responded to node counts later than the LIFG (see Supplementary Materials for converging evidence on the latency of the response based on analysis of the temporal derivative). This pattern of results, therefore, indicates that in production phrase-structure building operations preferentially took place before word onset in an incremental fashion. An analysis on the pausing patterns throughout the speech additionally revealed that top-down node counts affected pause length before word articulation, providing converging evidence for phrase-structure building to happen before word onset in production (see Supplementary Materials and Supplementary Fig. 3 for the analysis on pause length and word duration).

## Discussion

In the first study to investigate the neural correlates of syntactic processing during spontaneous production, we modelled incremental phrase-structure building with probabilistic parsers and used them to predict brain activity in BA44, BA45 and LpMTG. We found that phrase-structure operations successfully predicted brain activity during naturalistic speech. A central finding was that the timing of phrase-structure operations differed strongly between production and comprehension. The results suggest that phrase-structure building occurs in an integratory manner in comprehension, mostly in LpMTG and BA45. Phrase-structure building was instead markedly anticipatory in production (occurring predominantly before word onset), as evidenced by anticipatory parser operations predicting pause length before each word during speech, and by a highly anticipatory incremental production parser that best modelled the production data in BA45.

Syntactic processing elicited BOLD activity increases in both production and comprehension, but critically the temporal profiles of brain activity diverged across modalities. This highlights inherent processing differences between language production and comprehension. In production, structure building can proceed by establishing the upcoming structure before words are uttered, which was confirmed by the longer pauses associated with larger numbers of top-down parsing operations. In comprehension, instead, phrase-structure building proceeds in a more integratory manner, waiting for the input to commit to the structure. These results fit with previously obtained evidence on BOLD timing sensitivity to structure complexity and modality, where BOLD peaked earlier with more complex structures in production but later in comprehension, relative to easier structures (Giglio et al., 2022; Pallier et al., 2011; see Pylkkänen (2020; Pylkkänen et al., 2014) for converging evidence on production and comprehension dynamics of composition in MEG). Thus, the present results converge with previous controlled experiments in showing early syntactic encoding in production relative to later encoding in comprehension. This is likely due to different processing dynamics in production and comprehension, which have opposite inputs, outputs and mappings between linguistic levels (Pickering & Garrod, 2013).

It should be noted that these results only outline coarse processing dynamics, given the low temporal resolution of the BOLD signal, and that they do not aim to faithfully model all processes going on during speaking and listening. For example, these parsers are perfect ‘oracles’, meaning that they always posit phrase-structure building operations that actually happen, in contrast with potential ambiguities in the input (Hale, 2014). Recent evidence has shown that modelling syntactic ambiguity improves the fit with brain activity (Brennan et al., 2020). In addition, there is substantial evidence that comprehension is sensitive to the predictability of the input, such that some amount of anticipatory syntactic processing is expected in comprehension as well (Heilbron et al., 2022; Henderson et al., 2016; Shain et al., 2020; Willems et al., 2016). Indeed, Brennan et al. (2016) found a positive relationship between top-down operations, syntactic surprisal and BOLD activity in comprehension. Similarly, Coopmans et al. (*in prep*.) found that a top-down parser best modelled brain activity during comprehension in MEG. Nelson et al. (2017) instead found bottom-up counts to better model brain activity (measured with electrocorticography) than top-down counts for the comprehension of single sentences. It is possible that different characteristics of the speech input led to this difference between studies. In our case, the input was spontaneous speech that also included dysfluencies and corrections, while Brennan et al.’s and Coopmans at al.’s linguistic input were audiobook stories. There is evidence that lexical predictions can be influenced by reading strategies (Brothers et al., 2017). It might have been easier to anticipate the structure in the ‘cleaner’ audiobook story than in the recall of an unfamiliar story. The reduced contextual information available in Nelson et al. (2017) may also have led to a reduction in anticipatory syntactic processing. Future studies with naturalistic comprehension will need to clarify to what extent the nature of the input determines the strength of anticipatory vs. integratory syntactic structure building.

Previous studies found modality differences in the sensitivity of neural responses to syntactic processing (Giglio et al., 2022; Indefrey et al., 2004). In particular, syntactic processing led to stronger responses in production than comprehension. This difference could have been observed either due to task-related effects or due to modality-inherent differences, such as a stronger need in production to fully compute the syntactic structure to be able to speak correctly, in contrast to good-enough processing in comprehension (Bock, 1982; Ferreira et al., 2002; Garrett, 1980). While we could not directly address this question with modality as a between-subject variable, the results indicate that the different modality load on syntactic processing found in previous studies may in effect be task-related. In this study, syntactic structure building elicited a neural activity increase that was quantitatively similar across ROIs in both modalities, although with different dynamics. This finding provides evidence that, in contexts where production is spontaneous and unconstrained by artificial tasks and comprehension is meaningful and as a consequence more engaging, syntactic parsing and encoding have a similar load on brain activity.

Interestingly, there were some regional differences in the sensitivity to syntactic predictors in each modality. In particular, both LpMTG and BA45 responded to syntactic processing in comprehension. Instead, in production BA45 was the most responsive, with a less direct involvement of LpMTG activity (excluding a potential later activation as suggested by analysis of the temporal derivatives, see Supplementary Materials). This finding is somewhat unexpected, in light of previous results showing shared resources and representations across modalities (Giglio et al., 2022; Kempen et al., 2012; Segaert et al., 2012). It is not, however, the first study to show a higher comprehension load in temporal regions and a higher production load in frontal regions (Giglio et al., 2022; Humphreys & Gennari, 2014; Indefrey et al., 2004; Matchin & Wood, 2020). A plausible explanation relates to processing differences between modalities (Momma & Phillips, 2018). Both modalities may rely on the LIFG to coordinate syntactic processing (e.g. unification-type processing, Hagoort, 2013), while engaging the posterior temporal lobe for lexical-syntactic retrieval at different latencies depending on the amount of information available for structure building. In production, as suggested by the top-down parser, structure building may have been initiated before lexical-syntactic retrieval, engaging the LpMTG at later timescales, which is tentatively shown by the later response of the LpMTG in production. In comprehension, as indicated by the bottom-up parser, lexical-syntactic retrieval may have preceded or co-occurred with structure building and have appeared at canonical HRF delays. Interestingly, BA44 responded less strongly than BA45 to most predictors, in many cases with very low activity. This could suggest a reduced involvement of BA44 in syntactic processing in general, in line with Hagoort and Indefrey (2014) proposing a reduced function for BA44 in canonical syntactic processing. Alternatively, the anatomical selection may have been suboptimal for a region that is found to be highly variable between individuals (Fedorenko & Blank, 2020). Previous studies found also the anterior temporal lobe to show sensitivity to phrase-structure building operations (Brennan et al., 2012, 2016, 2020). We chose not to focus on this region due to converging evidence for a role of the anterior temporal lobe in semantic composition over syntactic processing (e.g. Lambon Ralph et al., 2017; Mesulam et al., 2014; Pylkkänen, 2020; Wilson et al., 2013), but we do not exclude that effects of phrase-structure building may have been present in the anterior temporal lobe in this study as well.

Returning to parser-specific modelling of syntactic processing, the parsers discussed so far were developed for syntactic processing specifically in comprehension (Hale, 2014). Since production is thought to proceed with different dynamics, we explored parsers that were more plausible for processing in production (Pickering & Garrod, 2013). In production, syntactic processing is thought to happen before word articulation (Bock & Levelt, 1994; Indefrey & Levelt, 2004; Levelt, 1989). There are different views on whether lexical access guides the structure, or whether the structure encoding the relations between concepts guides the order of lexical access (Bock & Ferreira, 2014). While the evidence provides mixed support for both accounts, suggesting that syntactic encoding is flexible and variable (e.g. Konopka & Meyer, 2014; Kuchinsky et al., 2011; van de Velde & Meyer, 2014), several proposals identify the verb as a central node in sentence planning, suggesting that the syntactic structure until the verb is sometimes computed early on (Levelt, 1989; Momma et al., 2016; Momma & Ferreira, 2019). Cross-linguistic evidence even suggests that in some languages some level of planning happens during the previous sentence (Sarvasy et al., 2022). By taking advantage of brain activity as an index of processing dynamics, we compared more and less incremental models of sentence planning with two parser models that made different predictions on the temporal unfolding of syntactic structure.

An incremental parser that is more anticipatory than the original top-down parser led to the best model fit, leading to the strongest increase in BA45 but not affecting LpMTG. Structure building thus proceeds before word articulation. This was also confirmed by converging results on longer pauses before words associated with more phrase-structure building operations, in line with previous behavioural evidence linking pausing patterns in speech with syntactic complexity (Ferreira, 1991). A less incremental parser that always plans the structures for a few chunks of words at a time provided the worst fit for brain activity. These results suggest that a highly incremental parser may be the more standard planning strategy in production, and that the structure up to the verb does not need to be planned at the start of the sentence. We thus provide the first piece of neuroimaging evidence addressing the long-standing debate on the incrementality of sentence planning. This approach could contribute to the understanding of the dynamics of sentence planning, by developing models that take into account the variability of each sentence, for example by modelling longer planning scopes only when the verb follows an internal argument (cf. Momma & Ferreira, 2019), or depending on word accessibility (Kuchinsky et al., 2011; van de Velde & Meyer, 2014).

Importantly, with this study we demonstrated the feasibility and benefits of studying production with spontaneous speech. The costs associated with spontaneous production, such as increased variability and dysfluencies of the linguistic signal, increased motion artifacts in fMRI and the slow temporal resolution, are balanced by the many advantages. Spontaneous production yields a larger amount of data than controlled tasks. This is the case especially in behavioural analysis but also with fMRI, provided the speech samples are of sufficient length. In addition, with spontaneous speech the artificiality of the task is largely reduced. Although speaking in monologue is not as common as dialogue, it is more ecologically valid than speaking following careful instructions with limited acceptable speech output. In addition, the probability distributions of linguistic inputs and outputs are preserved in spontaneous contexts, in contrast with many experiments (Jaeger, 2010). Finally, neuroimaging studies on spontaneous production allow for potentially new insight into production questions that have been so far mostly addressed with psycholinguistic studies.

In summary, we showed that spontaneous production can be used to study the neural correlates of linguistic processing, providing very rich data that can be directly linked to behavior with the analysis of pause length and word durations. We found that syntactic structure building engages the inferior frontal gyrus in both production and comprehension with diverging dynamics. Phrase-structure building was anticipatory in production but integratory in comprehension. Finally, we provided neural evidence for incremental models of syntactic encoding in production using production-specific parsers.

## Acknowledgements

We would like to thank Janice Chen and her lab for making the data available and especially for timestamping the recordings for word-by-word analysis; and Cas Coopmans for comments on an earlier version of the manuscript.

## Data availability

The fMRI data are available on OpenNeuro at the links provided in text. The word timestamps with linguistic annotations and the analysis code are available on request and will be made openly available upon publication.

## Materials and Methods

### Data acquisition and preprocessing

#### Production data

The production data used were collected by Chen et al. (2017) and made available on OpenNeuro (https://openneuro.org/datasets/ds001132/versions/1.0.0). In this experiment, participants watched an episode of the BBC TV series *Sherlock* and then recalled what happened in the episode. Data were collected for 22 right-handed native English participants (10 female, ages 18-26, mean age 20.8). Five participants were excluded due to excessive head motion (2 participants), because recall was shorter than 10 min (2 participants) or for falling asleep during the movie (1 participant). Data for one participant was not shared because of missing data at the end of the movie scan, which left us with 16 participants for the current analysis. Speaking led to an average framewise displacement of 0.32 (average per participant, range = 0.13-0.54), which was higher than the average in the comprehension data (0.22, range = 0.08-0.42) but was corrected for with noise regression (see fMRI data preprocessing for more details).

Participants watched the first 50 minutes of the first episode of the BBC TV series *Sherlock*, after confirming that they had not watched any episode of *Sherlock* before. Participants were told they would be asked to verbally describe what they had seen. After watching the episode, they were immediately instructed “to describe what they recalled of the movie in as much detail as they could, to try to recount events in the original order they were viewed in, and to speak for at least 10 min if possible but that longer was better. They were told that completeness and detail were more important than temporal order, and that if at any point they realized they had missed something, to return to it. Participants were then allowed to speak for as long as they wished, and verbally indicated when they were finished (for example, “I’m done”). During this session they were presented with a static black screen with a central white dot (but were not asked to, and did not, fixate).” Their speech was recorded in the scanner with an MR-compatible microphone.

We also used a second production scan for one of these participants, who also recalled an episode of BBC TV series *Merlin*, as part of the data collected and released by Zadbood et al. (2017). This speech sample was used as audio stimulus for the Comprehension data (see below). The procedure and acquisition were the same. Therefore, in total, we used 17 speech samples from 16 participants, since once participant recalled both *Sherlock* and *Merlin*. The 17 recalls were 10-45 minutes (mean = 22 minutes, SD = 8.8 minutes), including on average 2874 words (range = 1666-6230, SD = 1299).

#### Comprehension data

For the comprehension data, we used the data shared by Zadbood et al. (2017) on OpenNeuro (https://openneuro.org/datasets/ds001110/versions/00003). In this experiment, participants watched an episode of either BBC TV series *Merlin* or *Sherlock* and listened to an audio recording of the story they did not watch. Audio recordings were obtained from a participant that watched and recounted the two movies, here analysed as part of the production data. In this dataset, 52 right-handed native English speakers (age 18-45) were scanned. Fifteen participants were excluded because of head motion (n=4), for falling asleep (n=4), due to poor memory (n=5), for having seen the movie before (n=2). This resulted in 36 analysed participants, 18 that listened to the Merlin recall, and 18 that listened to the Sherlock recall. The audio recall for Merlin was 14.7 minutes long and included 2141 words. The audio recall for Sherlock was 17.5 minutes long and included 2468 words.

#### Data Acquisition

The acquisition parameters were identical in the two datasets. MRI data was collected on a 3T full-body scanner (Siemens Skyra) with a 20-channel head coil. Functional images were acquired using a T2*-weighted echo planar imaging pulse sequence (TR 1500 ms, TE 28 ms, flip angle 64, whole-brain coverage 27 slices of 4 mm thickness, in-plane resolution 3 × 3 mm^2^, FOV 192 × 192 mm^2^). Anatomical images were acquired using a T1-weighted MPRAGE pulse sequence (0.89 mm^3^ resolution).

#### fMRI data preprocessing

Preprocessing was performed using *fMRIPrep* 20.2.6 (Esteban, Markiewicz, et al. (2018); Esteban, Blair, et al. (2018)), which is based on *Nipype* 1.7.0 (Gorgolewski et al. (2011); Gorgolewski et al. (2018)).

##### Anatomical data preprocessing

The T1-weighted (T1w) image was corrected for intensity non-uniformity (INU) with N4BiasFieldCorrection (Tustison et al. 2010), distributed with ANTs 2.3.3 (Avants et al. 2008, RRID:SCR_004757), and used as T1w-reference throughout the workflow. The T1w-reference was then skull-stripped with a *Nipype* implementation of the antsBrainExtraction.sh workflow (from ANTs), using OASIS30ANTs as target template. Brain tissue segmentation of cerebrospinal fluid (CSF), white-matter (WM) and gray-matter (GM) was performed on the brain-extracted T1w using fast (FSL 5.0.9, RRID:SCR_002823, Zhang, Brady, and Smith 2001). Brain surfaces were reconstructed using recon-all (FreeSurfer 6.0.1, RRID:SCR_001847, Dale, Fischl, and Sereno 1999), and the brain mask estimated previously was refined with a custom variation of the method to reconcile ANTs-derived and FreeSurfer-derived segmentations of the cortical gray-matter of Mindboggle (RRID:SCR_002438, Klein et al. 2017).

##### Functional data preprocessing

For each BOLD run, the following preprocessing was performed. First, a reference volume and its skull-stripped version were generated using a custom methodology of *fMRIPrep*. Susceptibility distortion correction (SDC) was omitted, because no fieldmap was acquired. The BOLD reference was then co-registered to the T1w reference using bbregister (FreeSurfer) which implements boundary-based registration (Greve and Fischl 2009). Co-registration was configured with six degrees of freedom. Head-motion parameters with respect to the BOLD reference (transformation matrices, and six corresponding rotation and translation parameters) are estimated before any spatiotemporal filtering using mcflirt (FSL 5.0.9, Jenkinson et al. 2002). The BOLD time-series (including slice-timing correction when applied) were resampled onto their original, native space by applying the transforms to correct for head-motion. These resampled BOLD time-series will be referred to as *preprocessed BOLD in original space*, or just *preprocessed BOLD*. The BOLD time-series were resampled into standard space, generating a *preprocessed BOLD run in MNI152NLin2009cAsym space*. First, a reference volume and its skull-stripped version were generated using a custom methodology of *fMRIPrep*. Automatic removal of motion artifacts using independent component analysis (ICA-AROMA, Pruim et al. 2015) was performed on the *preprocessed BOLD on MNI space* time-series after removal of non-steady state volumes and spatial smoothing with an isotropic, Gaussian kernel of 6mm FWHM (full-width half-maximum). The “aggressive” noise-regressors were collected and placed in the corresponding confounds file. Several confounding time-series were calculated based on the *preprocessed BOLD:* framewise displacement (FD), the derivative of the relative (frame-to-frame) bulk head motion variance (DVARS) and three region-wise global signals. FD was computed using two formulations following Power (absolute sum of relative motions, Power et al. (2014)) and Jenkinson (relative root mean square displacement between affines, Jenkinson et al. (2002)). FD and DVARS are calculated for each functional run, both using their implementations in *Nipype* (following the definitions by Power et al. 2014). Additionally, a set of physiological regressors were extracted to allow for component-based noise correction (*CompCor*, Behzadi et al. 2007). Principal components are estimated after high-pass filtering the *preprocessed BOLD* time-series (using a discrete cosine filter with 128s cut-off) for anatomical CompCor (aCompCor). For aCompCor, three probabilistic masks (cerebrospinal fluid (CSF), white matter (WM) and combined CSF+WM) are generated in anatomical space.

### Incremental complexity metrics

#### Phrase-structure parsing

First, we extracted the constituent structure of each sentence with a probabilistic context-free phrase-structure grammar. We used the Stanford parser with CoreNLP in Python 3 via the Natural Language Toolkit (NLTK) package (Klein & Manning, 2003; Manning et al., 2014). The transcript provided in the shared dataset was divided in what we considered independent sentences. Since the production was very spontaneous and unconstrained, sentence boundaries were not objective and self-evident as they are in text. In speech, the boundaries can depend on the syntactic structure of the sentence, but also on pausing patterns. For example, coordinated clauses may be considered one single sentence or divided into two separate sentences based on pause lengths. After extensive exposure to the transcripts, it became clear that shorter boundaries better reflect the planning chunks followed by speakers. In particular, some sentences extend over 30 words or more, with many embedded phrases. Participants, however, do not appear to fully keep in working memory the original syntactic structure, which is revealed by their dysfluencies and corrections throughout the long sentence. Dysfluencies affect how sentence boundaries, which are critical for the performance of automatized parsers, can be placed in the discourse, since they are not as obvious as in text. For example, boundaries could fully track the syntactic structure, also including hesitations and corrections within its boundaries, or they could track speech patterns and ‘reset’ every time there is a dysfluency. In this type of data, with monologues without audience feedback, short dysfluency boundaries seemed more appropriate, but it is to be determined if different approaches work better in other contexts. It should be noted that an initial analysis was run on more liberal sentences, which perhaps better tracked the overarching syntactic structure but did not optimally reflect the planning processes of participants. The results were similar with both sentence boundaries approaches, but the conservative approach to sentence length was less noisy.

Following sentence parsing with the Stanford constituent parser, we took a measure of syntactic processing with incremental complexity metrics derived from the number of syntactic nodes that are built with each word. Nodes can be built with different parsing strategies: top-down, bottom-up and left-corner (Hale, 2014). In top-down parsing, nodes are built from the top of the syntactic tree to the terminal node (corresponding to a word). In other words, nodes are counted when phrases are opened. This strategy can lead to the anticipation of nodes that may not always be known to a listener. For example, in the sentence “Mary eats apples daily”, a node accounting for the upcoming presence of “daily” is counted already at the word “eats”. This anticipation is justifiable in production, where the upcoming structure is presumably known to the speaker in advance, but it might reflect unjustifiable prediction in comprehension. Nevertheless, this implementation of a top-down strategy may be successful in accounting for predictive processes in comprehension.

At the other end of the incremental parsing spectrum is bottom-up parsing, according to which nodes are built from the bottom of the syntactic tree (i.e. from the terminal nodes, corresponding to each word) up to the highest *closed* nodes, i.e. nodes where all daughter nodes have already been met. For example, in Figure 1B, the top node in purple (S) cannot be built until its right-branching node VP is built as well, which in this case only happens at the end of the sentence. In other words, bottom-up parsing builds nodes when phrases are closed. This strategy thus predicts increased syntactic processing at the end of clauses and sentences, after all the evidence for the structure is encountered. We expected this parsing strategy to better reflect processing in comprehension than production, because in the latter the structure is presumably already built before the last word is uttered. Neither top-down nor bottom-up parsing strategies fully match human performance (Brennan & Pylkkänen, 2017; Hale, 2014), but they capture aspects of syntactic processing that are expected to differ across modalities. Finally, left-corner parsing needs less evidence than bottom-up parsing to count nodes, but is not as predictive as top-down parsing. After convolving with the haemodynamic response function, left-corner was highly correlated with the top-down parser (Supplementary Fig. 4). Therefore, we decided to only focus on opposite parsing strategies that were most expected to differ between production and comprehension, i.e. top-down and bottom-up.

We also counted the number of nodes that were still open at each word with an *open nodes* measure, similarly to Nelson et al., (2017). Open nodes were the number of nodes that were open at each word: this measure tracked the number of nodes that had been opened up to the word and that had not been closed yet, thus providing an index for the number of nodes that need to be kept in working memory until they can be merged in a constituent (Nelson et al., 2017).

#### Production-specific parsing operations

To account for the timing that is specific to production, we developed two production-specific parsers. An *early top-down* model counts the nodes that are built for the *next* word. At the first word of the sentence, nodes are counted for the first and second word (even though nodes built for the first word would have been built earlier, we preferred this over making assumptions on *when* the nodes would be built before the sentence, which could be varying due to different factors). At the second word, nodes are counted for the third word, etc.

For the less-incremental *chunked* parsing, we selected the heads of each sentence following dependency parsing (see Supplementary Materials for more information on the analysis on dependency parsing). We considered as heads all words that had a dependent relation attached to them (e.g. the verb is head of subject and object). We then counted all nodes (of the same constituent structure used by the other parsers) encountered from the first word up to and including the next head, then from the head up to and including the next head, and so on. Chunked parsing, therefore, builds nodes early on for all the upcoming words that are dependent relations of the next head. For example, at the start of a sentence all the nodes are built for the structure up to and including the verb, usually the first head.

It should be noted that *top-down, early top-down*, and *chunked* measures were highly correlated after convolving with the haemodynamic response function (Supplementary Fig. 4). To avoid collinearity, instead of comparing them in the same model, we tested models with only one predictor and determined which model provided the best fit (see Regression analysis for more details).

#### Word surprisal

We quantified word surprisal from transformer model GPT-2 (Radford et al., 2019). We used GPT-2 XL via the TensorFlow implementation provided by HuggingFace’s Transformers package (Wolf et al., 2020). Each word’s probability was based on a context of at least 700 words after the first 700 words of each participant’s recall. Surprisal was calculated as the negative logarithm of the conditional probability of the word based on context.

### Behavioural analysis

To determine if these indexes of processing complexity had an effect on participants’ speech patterns, we inspected how they affected word duration and pause lengths in all the production recalls. Recordings were not made available with the Production dataset, but word timestamps for each participant’s recall were shared by Janice Chen’s lab. Onsets and offsets of each word were obtained with *Gentle* (https://lowerquality.com/gentle/, https://github.com/lowerquality/gentle). We ran a linear mixed-effects model with *lme4* (version 1.1-26, Bates et al., 2015) in R (version 4.0.3). We used number of syllables, word frequency, word surprisal, top-down, bottom-up and open nodes as predictors for pause length (before the word characterized by each predictor) and word duration. This analysis allowed us to compare neural effects with behavioural patterns of speech.

### fMRI analysis

#### Predictor timeseries

Each word-by-word predictor was mean-centered (except for the word rate predictor) and convolved with the canonical haemodynamic response function following SPM’s double gamma function as computed in *nilearn*. We thus obtained predictor timeseries temporally resampled to the acquisition TR of 1.5 s, reflecting BOLD increases and decreases following predictor weights time-locked to word onset (Fig. 2C).

#### ROI selection

We selected 3 ROIs that have been associated with syntactic processing in previous studies: two LIFG ROIs, following the distinction between LIFG *pars opercularis* (BA44) and LIFG *pars triangularis* (BA45), and left posterior middle temporal gyrus (LpMTG). After preprocessing the fMRI data, we selected the ROIs for each participant in their functional space. BA44 and BA45 were extracted following Freesurfer’s label creation with the Destrieux Atlas (Destrieux et al., 2010) and resampled to functional space with bbregister. Freesurfer’s MTG ROI is quite long in extension, following the gyrus from very posterior portions to the temporal pole. We therefore extracted this ROI and then masked it with a posterior temporal lobe mask (posterior to Heschl’s gyrus) based on the Harvard-Oxford cortical atlas. Examples of these ROIs in MNI brain can be seen in Figure 2E.

#### Timeseries extraction

The BOLD timeseries were extracted with NiftiLabelsMasker from nilearn (Abraham et al., 2014; *Nilearn/Nilearn*, 2011/2022), after confound regression. Framewise displacement, DVARS, motion parameters, aCompCor paramers and ICA-AROMA regressors classified as noise were used for noise regression, to reduce the impact of motion artifacts caused by speaking. The timeseries was extracted from the functional BOLD volumes in functional space as an average of the voxels in each ROI mask.

#### Regression analysis

To determine to what extent each of these continuous indices of syntactic processing significantly affected brain activity (average BOLD activity in the three ROIs), we used linear mixed-effects models with *lme4* (version 1.1-26, Bates et al., 2015) in R (version 4.0.3). We used a baseline model that included word rate (i.e. a predictor indicating the onset of each word), syllable rate, as an index of articulatory rate, log-transformed word frequency, and word surprisal. All models additionally included modality and ROI as factors. Modality (production vs. comprehension) was contrast-coded with deviation coding. We used Helmert coding for ROI, contrasting LIFG with LpMTG, and the two LIFG *partes* with each other. All other factors were continuous numerical predictors. All models included word surprisal and its interaction with ROI and modality. All models also included by-participant random slopes for syllable rate, frequency, word surprisal and other factors of interest, excluding by-participant random effects and correlations to allow for convergence and avoid singularity issues. In some cases, we had to exclude the random slopes for one of these factors, but never for the factor of interest in that model. We computed the contribution of factors to the models using *car* (version 3.0-10, Fox et al., 2021), and pairwise comparisons with the package *emmeans* (version 1.6.1, Lenth et al., 2022).

The first model determined the contribution of top-down and bottom-up metrics of phrase-structure building to brain activity in the three ROIs and in each modality to a baseline model that included word surprisal and open nodes. The interactions of each metric with ROI and modality were included and the significant contribution of the incremental metric in a region or modality was determined with pairwise comparisons. With this model we also determined to what extent word surprisal and open nodes affected brain activity in each modality.

We then used three models to ask whether metrics of syntactic processing fine-tuned for production would improve model fit. These metrics are not realistic for syntactic processing in comprehension, so the models only included production data. The baseline models all included word surprisal and bottom-up parser operations, and additionally included *top-down*, or *early top-down*, or *chunked* predictors of phrase structure building and their relative by-participant random slopes. Since the three parsers were highly correlated after convolving with the HRF, we separately fitted three linear models. We compared model fit with the Akaike Information Criterion (AIC), where more negative values indicate better model fit (Cavanaugh & Neath, 2019).

## References

Abraham, A., Pedregosa, F., Eickenberg, M., Gervais, P., Mueller, A., Kossaifi, J., Gramfort, A., Thirion, B., & Varoquaux, G. (2014). Machine learning for neuroimaging with scikit-learn. Frontiers in Neuroinformatics, 8. https://doi.org/10.3389/fninf.2014.00014

Alday, P. M., Schlesewsky, M., & Bornkessel-Schlesewsky, I. (2017). Electrophysiology Reveals the Neural Dynamics of Naturalistic Auditory Language Processing: Event-Related Potentials Reflect Continuous Model Updates. ENeuro, 4(6). https://doi.org/10.1523/ENEURO.0311-16.2017

Andric, M., & Small, S. L. (2015). FMRI methods for studying the neurobiology of language under naturalistic conditions. In R. M. Willems (Ed.), Cognitive Neuroscience of Natural Language Use (pp. 8–28). Cambridge University Press. https://doi.org/10.1017/CBO9781107323667.002

Bates, D., Mächler, M., Bolker, B., & Walker, S. (2015). Fitting Linear Mixed-Effects Models Using lme4. Journal of Statistical Software, 67(1), 1–48. https://doi.org/10.18637/jss.v067.i01

Bhattasali, S., Fabre, M., Luh, W.-M., Saied, H. A., Constant, M., Pallier, C., Brennan, J. R., Spreng, R. N., & Hale, J. (2019). Localising memory retrieval and syntactic composition: An fMRI study of naturalistic language comprehension. Language, Cognition and Neuroscience, 34(4), 491–510. https://doi.org/10.1080/23273798.2018.1518533

Bock, K. (1982). Toward a cognitive psychology of syntax: Information processing contributions to sentence formulation. Psychological Review, 89(1), 1–47. https://doi.org/10.1037/0033-295X.89.1.1

Bock, K., & Ferreira, V. (2014). Syntactically speaking. In The Oxford handbook of language production (pp. 21–46). Oxford University Press. https://doi.org/10.1093/oxfordhb/9780199735471.013.008

Bock, K., & Levelt, W. J. M. (1994). Language production: Grammatical encoding. Handbook of psycholinguistics. ed. by Morton A. Gernsbacher, 945–984. San Diego, CA: Academic Press.

Brennan, J. R. (2016). Naturalistic Sentence Comprehension in the Brain. Language and Linguistics Compass, 10(7), 299–313. https://doi.org/10.1111/lnc3.12198

Brennan, J. R., Dyer, C., Kuncoro, A., & Hale, J. T. (2020). Localizing syntactic predictions using recurrent neural network grammars. Neuropsychologia, 146, 107479. https://doi.org/10.1016/j.neuropsychologia.2020.107479

Brennan, J. R., & Hale, J. T. (2019). Hierarchical structure guides rapid linguistic predictions during naturalistic listening. PLOS ONE, 14(1), e0207741. https://doi.org/10.1371/journal.pone.0207741

Brennan, J. R., Nir, Y., Hasson, U., Malach, R., Heeger, D. J., & Pylkkänen, L. (2012). Syntactic structure building in the anterior temporal lobe during natural story listening. Brain and Language, 120(2), 163–173. https://doi.org/10.1016/j.bandl.2010.04.002

Brennan, J. R., & Pylkkänen, L. (2017). MEG Evidence for Incremental Sentence Composition in the Anterior Temporal Lobe. Cognitive Science, 41(S6), 1515–1531. https://doi.org/10.1111/cogs.12445

Brennan, J. R., Stabler, E. P., Van Wagenen, S. E., Luh, W.-M., & Hale, J. T. (2016). Abstract linguistic structure correlates with temporal activity during naturalistic comprehension. Brain and Language, 157–158, 81–94. https://doi.org/10.1016/j.bandl.2016.04.008

Brothers, T., Swaab, T. Y., & Traxler, M. J. (2017). Goals and strategies influence lexical prediction during sentence comprehension. Journal of Memory and Language, 93, 203–216. https://doi.org/10.1016/j.jml.2016.10.002

Cavanaugh, J. E., & Neath, A. A. (2019). The Akaike information criterion: Background, derivation, properties, application, interpretation, and refinements. WIREs Computational Statistics, 11(3), e1460. https://doi.org/10.1002/wics.1460

Destrieux, C., Fischl, B., Dale, A., & Halgren, E. (2010). Automatic parcellation of human cortical gyri and sulci using standard anatomical nomenclature. NeuroImage, 53(1), 1–15. https://doi.org/10.1016/j.neuroimage.2010.06.010

Fedorenko, E., & Blank, I. A. (2020). Broca’s Area Is Not a Natural Kind. Trends in Cognitive Sciences, 24(4), 270–284. https://doi.org/10.1016/j.tics.2020.01.001

Ferreira, F. (1991). Effects of length and syntactic complexity on initiation times for prepared utterances. Journal of Memory and Language, 30(2), 210–233.

Ferreira, F. (2003). The misinterpretation of noncanonical sentences. Cognitive Psychology, 47(2), 164–203. https://doi.org/10.1016/S0010-0285(03)00005-7

Ferreira, F., Bailey, K. G. D., & Ferraro, V. (2002). Good-Enough Representations in Language Comprehension. Current Directions in Psychological Science, 11(1), 11–15. https://doi.org/10.1111/1467-8721.00158

Ferreira, F., & Lowder, M. W. (2016). Chapter Six—Prediction, Information Structure, and Good-Enough Language Processing. In B. H. Ross (Ed.), Psychology of Learning and Motivation (Vol. 65, pp. 217–247). Academic Press. https://doi.org/10.1016/bs.plm.2016.04.002

Fox, J., Weisberg, S., Price, B., Adler, D., Bates, D., Baud-Bovy, G., Bolker, B., Ellison, S., Firth, D., Friendly, M., Gorjanc, G., Graves, S., Heiberger, R., Krivitsky, P., Laboissiere, R., Maechler, M., Monette, G., Murdoch, D., Nilsson, H., … R-Core. (2021). car: Companion to Applied Regression (3.0-12). https://CRAN.R-project.org/package=car

Garrett, M. F. (1980). Levels of processing in sentence production. In Language production Vol. 1: Speech and talk (pp. 177–220). Academic Press.

Garrett, M. F. (1982). Remarks on the relation between language production and language comprehension systems. In Neural models of language processes (pp. 209–224). Elsevier.

Giglio, L., Ostarek, M., Weber, K., & Hagoort, P. (2022). Commonalities and Asymmetries in the Neurobiological Infrastructure for Language Production and Comprehension. Cerebral Cortex, 32(7), 1405–1418. https://doi.org/10.1093/cercor/bhab287

Goldberg, A. E., & Ferreira, F. (2022). Good-enough language production. Trends in Cognitive Sciences, 26(4), 300–311. https://doi.org/10.1016/j.tics.2022.01.005

Griffin, Z. M., & Bock, K. (2000). What the Eyes Say About Speaking: Psychological Science. https://journals.sagepub.com/doi/10.1111/1467-9280.00255

Hagoort, P. (2013). MUC (Memory, Unification, Control) and beyond. Frontiers in Psychology, 4, 416. https://doi.org/10.3389/fpsyg.2013.00416

Hagoort, P., & Indefrey, P. (2014). The Neurobiology of Language Beyond Single Words. Annual Review of Neuroscience, 37(1), 347–362. https://doi.org/10.1146/annurev-neuro-071013-013847

Hale, J. T. (2014). Automaton theories of human sentence comprehension. Center for the Study of Language and Information.

Hale, J. T., Campanelli, L., Li, J., Bhattasali, S., Pallier, C., & Brennan, J. R. (2022). Neurocomputational Models of Language Processing. Annual Review of Linguistics, 8(1), 427–446. https://doi.org/10.1146/annurev-linguistics-051421-020803

Heilbron, M., Armeni, K., Schoffelen, J.-M., Hagoort, P., & de Lange, F. P. (2022). A hierarchy of linguistic predictions during natural language comprehension. Proceedings of the National Academy of Sciences, 119(32), e2201968119. https://doi.org/10.1073/pnas.2201968119

Henderson, J. M., Choi, W., Lowder, M. W., & Ferreira, F. (2016). Language structure in the brain: A fixation-related fMRI study of syntactic surprisal in reading. NeuroImage, 132, 293–300. https://doi.org/10.1016/j.neuroimage.2016.02.050

Hu, J., Small, H., Kean, H., Takahashi, A., Zekelman, L., Kleinman, D., Ryan, E., Nieto-Castañón, A., Ferreira, V., & Fedorenko, E. (2022). Precision fMRI reveals that the language-selective network supports both phrase-structure building and lexical access during language production. Cerebral Cortex, bhac350. https://doi.org/10.1093/cercor/bhac350

Huizeling, E., Peeters, D., & Hagoort, P. (2021). Prediction of upcoming speech under fluent and disfluent conditions: Eye tracking evidence from immersive virtual reality. Language, Cognition and Neuroscience, 0(0), 1–28. https://doi.org/10.1080/23273798.2021.1994621

Humphreys, G. F., & Gennari, S. P. (2014). Competitive mechanisms in sentence processing: Common and distinct production and reading comprehension networks linked to the prefrontal cortex. NeuroImage, 84, 354–366. https://doi.org/10.1016/j.neuroimage.2013.08.059

Indefrey, P. (2011). The Spatial and Temporal Signatures of Word Production Components: A Critical Update. Frontiers in Psychology, 2. https://doi.org/10.3389/fpsyg.2011.00255

Indefrey, P. (2018). The Relationship Between Syntactic Production and Comprehension. The Oxford Handbook of Psycholinguistics. https://doi.org/10.1093/oxfordhb/9780198786825.013.20

Indefrey, P., Hellwig, F., Herzog, H., Seitz, R. J., & Hagoort, P. (2004). Neural responses to the production and comprehension of syntax in identical utterances. Brain and Language, 89(2), 312–319. https://doi.org/10.1016/S0093-934X(03)00352-3

Indefrey, P., & Levelt, W. J. M. (2004). The spatial and temporal signatures of word production components. Cognition, 92(1), 101–144. https://doi.org/10.1016/j.cognition.2002.06.001

Jaeger, T. F. (2010). Redundancy and reduction: Speakers manage syntactic information density. Cognitive Psychology, 61(1), 23–62. https://doi.org/10.1016/j.cogpsych.2010.02.002

Kempen, G., Olsthoorn, N., & Sprenger, S. (2012). Grammatical workspace sharing during language production and language comprehension: Evidence from grammatical multitasking. Language and Cognitive Processes, 27(3), 345–380. https://doi.org/10.1080/01690965.2010.544583

Klein, D., & Manning, C. D. (2003). Accurate unlexicalized parsing. Proceedings of the 41st Annual Meeting on Association for Computational Linguistics - ACL ‘03, 1, 423–430. https://doi.org/10.3115/1075096.1075150

Konopka, A. E., & Meyer, A. S. (2014). Priming sentence planning. Cognitive Psychology, 73, 1–40. https://doi.org/10.1016/j.cogpsych.2014.04.001

Kuchinsky, S. E., Bock, K., & Irwin, D. E. (2011). Reversing the hands of time: Changing the mapping from seeing to saying. Journal of Experimental Psychology: Learning, Memory, and Cognition, 37(3), 748–756. https://doi.org/10.1037/a0022637

Lambon Ralph, M. A., Jefferies, E., Patterson, K., & Rogers, T. T. (2017). The neural and computational bases of semantic cognition. Nature Reviews Neuroscience, 18(1), 42–55. https://doi.org/10.1038/nrn.2016.150

Lenth, R. V., Buerkner, P., Herve, M., Love, J., Miguez, F., Riebl, H., & Singmann, H. (2022). emmeans: Estimated Marginal Means, aka Least-Squares Means (1.7.3). https://CRAN.R-project.org/package=emmeans

Levelt, W. J. M. (1989). Speaking: From intention to articulation. ACL. MIT Press Series in Natural-Language Processing. MIT Press, Cambridge, Massachusetts.

Li, J., & Hale, J. (2019). Grammatical predictors for fMRI timecourses. Minimalist Parsing, 159–173.

Lopopolo, A., van den Bosch, A., Petersson, K.-M., & Willems, R. M. (2021). Distinguishing Syntactic Operations in the Brain: Dependency and Phrase-Structure Parsing. Neurobiology of Language, 2(1), 152–175. https://doi.org/10.1162/nol_a_00029

Manning, C., Surdeanu, M., Bauer, J., Finkel, J., Bethard, S., & McClosky, D. (2014). The Stanford CoreNLP Natural Language Processing Toolkit. Proceedings of 52nd Annual Meeting of the Association for Computational Linguistics: System Demonstrations, 55–60. https://doi.org/10.3115/v1/P14-5010

Matchin, W., & Hickok, G. (2016). ‘Syntactic Perturbation’ During Production Activates the Right IFG, but not Broca’s Area or the ATL. Frontiers in Psychology, 7.https://doi.org/10.3389/fpsyg.2016.00241

Matchin, W., & Wood, E. (2020). Syntax-sensitive regions of the posterior inferior frontal gyrus and the posterior temporal lobe are differentially recruited by production and perception. Cerebral Cortex Communications. https://doi.org/10.1093/texcom/tgaa029

Menenti, L., Gierhan, S. M. E., Segaert, K., & Hagoort, P. (2011). Shared Language: Overlap and Segregation of the Neuronal Infrastructure for Speaking and Listening Revealed by Functional MRI. Psychological Science, 22(9), 1173–1182. https://doi.org/10.1177/0956797611418347

Mesulam, M.-M., Rogalski, E. J., Wieneke, C., Hurley, R. S., Geula, C., Bigio, E. H., Thompson, C. K., & Weintraub, S. (2014). Primary progressive aphasia and the evolving neurology of the language network. Nature Reviews Neurology, 10(10), 554–569. https://doi.org/10.1038/nrneurol.2014.159

Momma, S., & Ferreira, V. S. (2019). Beyond linear order: The role of argument structure in speaking. Cognitive Psychology, 114, 101228. https://doi.org/10.1016/j.cogpsych.2019.101228

Momma, S., & Phillips, C. (2018). The Relationship Between Parsing and Generation. Annual Review of Linguistics, 4(1), 233–254. https://doi.org/10.1146/annurev-linguistics-011817-045719

Momma, S., Slevc, L. R., & Phillips, C. (2016). The timing of verb selection in Japanese sentence production. Journal of Experimental Psychology: Learning, Memory, and Cognition, 42(5), 813–824. https://doi.org/10.1037/xlm0000195

Nelson, M. J., Karoui, I. E., Giber, K., Yang, X., Cohen, L., Koopman, H., Cash, S. S., Naccache, L., Hale, J. T., Pallier, C., & Dehaene, S. (2017). Neurophysiological dynamics of phrase-structure building during sentence processing. Proceedings of the National Academy of Sciences, 114(18), E3669–E3678. https://doi.org/10.1073/pnas.1701590114 Nilearn/nilearn. (2022). [Python]. nilearn. https://github.com/nilearn/nilearn (Original work published 2011)

Pallier, C., Devauchelle, A.-D., & Dehaene, S. (2011). Cortical representation of the constituent structure of sentences. Proceedings of the National Academy of Sciences, 108(6), 2522–2527. https://doi.org/10.1073/pnas.1018711108

Peeters, D. (2019). Virtual reality: A game-changing method for the language sciences. Psychonomic Bulletin & Review, 26(3), 894–900. https://doi.org/10.3758/s13423-019-01571-3

Pickering, M. J., & Garrod, S. (2013). An integrated theory of language production and comprehension. Behavioral and Brain Sciences, 36(4), 329–347. https://doi.org/10.1017/S0140525X12001495

Pylkkänen, L. (2020). Neural basis of basic composition: What we have learned from the red–boat studies and their extensions. Philosophical Transactions of the Royal Society B: Biological Sciences, 375(1791), 20190299. https://doi.org/10.1098/rstb.2019.0299

Pylkkänen, L., Bemis, D. K., & Blanco Elorrieta, E. (2014). Building phrases in language production: An MEG study of simple composition. Cognition, 133(2), 371–384. https://doi.org/10.1016/j.cognition.2014.07.001

Radford, A., Wu, J., Child, R., Luan, D., Amodei, D., & Sutskever, I. (n.d.). Language Models are Unsupervised Multitask Learners. 24.

Sarvasy, H. S., Morgan, A. M., Yu, J., Ferreira, V. S., & Momma, S. (2022). Cross-clause planning in Nungon (Papua New Guinea): Eye-tracking evidence. Memory & Cognition. https://doi.org/10.3758/s13421-021-01253-3

Segaert, K., Menenti, L., Weber, K., Petersson, K. M., & Hagoort, P. (2012). Shared Syntax in Language Production and Language Comprehension—An fMRI Study. Cerebral Cortex (New York, NY), 22(7), 1662–1670. https://doi.org/10.1093/cercor/bhr249

Shain, C., Blank, I.A., van Schijndel, M., Schuler, W., & Fedorenko, E. (2020). FMRI reveals language-specific predictive coding during naturalistic sentence comprehension. Neuropsychologia, 138, 107307. https://doi.org/10.1016/j.neuropsychologia.2019.107307

Snijders, T. M., Vosse, T., Kempen, G., Van Berkum, J. J. A., Petersson, K. M., & Hagoort, P. (2009). Retrieval and Unification of Syntactic Structure in Sentence Comprehension: An fMRI Study Using Word-Category Ambiguity. Cerebral Cortex, 19(7), 1493–1503. https://doi.org/10.1093/cercor/bhn187

Stanojević, M., Bhattasali, S., Dunagan, D., Campanelli, L., Steedman, M., Brennan, J., & Hale, J. (2021). Modeling Incremental Language Comprehension in the Brain with Combinatory Categorial Grammar. Proceedings of the Workshop on Cognitive Modeling and Computational Linguistics, 23–38. https://doi.org/10.18653/v1/2021.cmcl-1.3

Takashima, A., Konopka, A., Meyer, A., Hagoort, P., & Weber, K. (2020). Speaking in the Brain: The Interaction between Words and Syntax in Sentence Production. Journal of Cognitive Neuroscience, 32(8), 1466–1483. https://doi.org/10.1162/jocn_a_01563

Uddén, J., Hultén, A., Schoffelen, J.-M., Lam, N., Harbusch, K., Bosch, A. van den, Kempen, G., Petersson, K. M., & Hagoort, P. (2019). Supramodal Sentence Processing in the Human Brain: Fmri Evidence for the Influence of Syntactic Complexity in More Than 200 Participants. BioRxiv, 576769. https://doi.org/10.1101/576769

van de Velde, M., & Meyer, A. S. (2014). Syntactic flexibility and planning scope: The effect of verb bias on advance planning during sentence recall. Frontiers in Psychology, 5. https://doi.org/10.3389/fpsyg.2014.01174

Wehbe, L., Murphy, B., Talukdar, P., Fyshe, A., Ramdas, A., & Mitchell, T. (2014). Simultaneously Uncovering the Patterns of Brain Regions Involved in Different Story Reading Subprocesses. PLOS ONE, 9(11), e112575. https://doi.org/10.1371/journal.pone.0112575

Willems, R. M., Frank, S. L., Nijhof, A. D., Hagoort, P., & van den Bosch, A. (2016). Prediction During Natural Language Comprehension. Cerebral Cortex, 26(6), 2506–2516. https://doi.org/10.1093/cercor/bhv075

Willems, R. M., & Gerven, M. A. J. van. (2018). New FMRI Methods for the Study of Language. The Oxford Handbook of Psycholinguistics. https://doi.org/10.1093/oxfordhb/9780198786825.013.42

Wilson, S. M., DeMarco, A. T., Henry, M. L., Gesierich, B., Babiak, M., Mandelli, M. L., Miller, B. L., & Gorno-Tempini, M. L. (2013). What Role Does the Anterior Temporal Lobe Play in Sentence-level Processing? Neural Correlates of Syntactic Processing in Semantic Variant Primary Progressive Aphasia. Journal of Cognitive Neuroscience, 26(5), 970–985. https://doi.org/10.1162/jocn_a_00550

Wolf, T., Debut, L., Sanh, V., Chaumond, J., Delangue, C., Moi, A., Cistac, P., Rault, T., Louf, R., Funtowicz, M., Davison, J., Shleifer, S., von Platen, P., Ma, C., Jernite, Y., Plu, J., Xu, C., Le Scao, T., Gugger, S., … Rush, A. (2020). Transformers: State-of-the-Art Natural Language Processing. Proceedings of the 2020 Conference on Empirical Methods in Natural Language Processing: System Demonstrations, 38–45. https://doi.org/10.18653/v1/2020.emnlp-demos.6

Zaccarella, E., Meyer, L., Makuuchi, M., & Friederici, A. D. (2017). Building by Syntax: The Neural Basis of Minimal Linguistic Structures. Cerebral Cortex, 27(1), 411–421. https://doi.org/10.1093/cercor/bhv234

Zaccarella, E., Schell, M., & Friederici, A. D. (2017). Reviewing the functional basis of the syntactic Merge mechanism for language: A coordinate-based activation likelihood estimation meta-analysis. Neuroscience & Biobehavioral Reviews, 80, 646–656. https://doi.org/10.1016/j.neubiorev.2017.06.011

