## Supplementary Materials for "Diverging Neural Dynamics for Syntactic Structure Building in Naturalistic Speaking and Listening"

### Dependency parsing

We extracted dependency parsing as an index to perform chunking of each sentence in relational units. We used the dependency parser provided by the Stanford parser via CoreNLP. We identified the heads of the sentence as the words that have a relation attached to them. For example, in the sentence “He’s examining one of the bodies”, “examining” is the first head, followed by “one” and by “bodies” (Supplementary Fig. 1). These three words are the only ones that have a dependency relation attached to them, while the other words are all dependents of one head. The *chunked* parsing strategy counted the nodes intervening between all heads, to model a less incremental strategy to syntactic structure building, following the idea that speakers plan the structure of a few words at a time (e.g. always planning the structure of the verb at the start of the sentence).

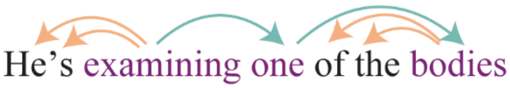


**Supplementary Figure 1**: *Dependency parse of the sentence. Left-relations are in orange, right relations in green, heads are in purple. Heads are words on which a dependency relation is attached (i.e. from which an arrow starts).*

### Temporal derivative

Since the results of production-specific parsers indicated that the LpMTG may have had a later response to top-down operations than BA45, we looked into how the temporal derivatives of the same predictors modelled the data. The temporal derivative of the haemodynamic response function (HRF) is usually used in fMRI analysis to account for small differences in the latency of the BOLD response. An increase in the temporal derivatives means that the BOLD response peaks earlier, while a decrease indicates a later peak. We ran the same linear-mixed effects model used before with the addition of temporal derivatives of all predictors of interest (Supplementary Table 5). We found a significant three-way interaction between the top-down derivative, modality and ROI ($\chi^2 = 15.7$, $p < 0.0004$); the surprisal derivative, modality and ROI ($\chi^2 = 18$, $p < 0.0002$); and two-way interactions between the open nodes derivative and modality ($\chi^2 = 4.4$, $p < 0.037$), and the open nodes derivative and ROI ($\chi^2 = 26.8$, $p < 0.0001$). The top-down temporal derivative was marginally significantly positive in BA45 in production (estimate = 0.31, SE = 0.16, $p = 0.056$), indicating an earlier BOLD peak than assumed by the canonical HRF, while it was significantly negative in LpMTG (estimate = 0.32, SE = 0.16, $p = 0.046$), indicating a later peak. It was not significantly different from zero in comprehension, nor did it differ between ROIs. These results suggest that the LpMTG may have been active after BA45 in response to more top-down node counts.

Word surprisal elicited later BOLD peaks in comprehension and earlier BOLD peaks in production, relative to the canonical HRF. In comprehension, BA45 and LpMTG were both significantly related to a decrease in activity (BA45: estimate = 0.46, SE = 0.16, $p = 0.003$; LpMTG: estimate = 0.55, SE = 0.16, $p = 0.0005$). In production, both activity in both BA44 and BA45 increased with the temporal derivative for surprisal (BA44: estimate = 0.53, SE = 0.2, $p = 0.16$; BA45: estimate = 0.97, SE = 0.2, $p < 0.0001$). These results suggest that word surprisal elicited earlier activity increases in production than comprehension, which likely relates to the timing of lexical access (before word onset in production, after word onset in comprehension). The open nodes measure showed earlier BOLD responses in the

LpMTG (estimate = 1.3, SE = 0.3, $p < 0.0001$), and earlier responses in comprehension than production (estimate = 0.213, SE =0.1, $p = 0.037$). Again, BA45 and the LpMTG had different BOLD peak latencies, suggesting that BA45 responded earlier to top-down nodes but later to open nodes, while LpMTG responded earlier to open nodes and later to top-down nodes.

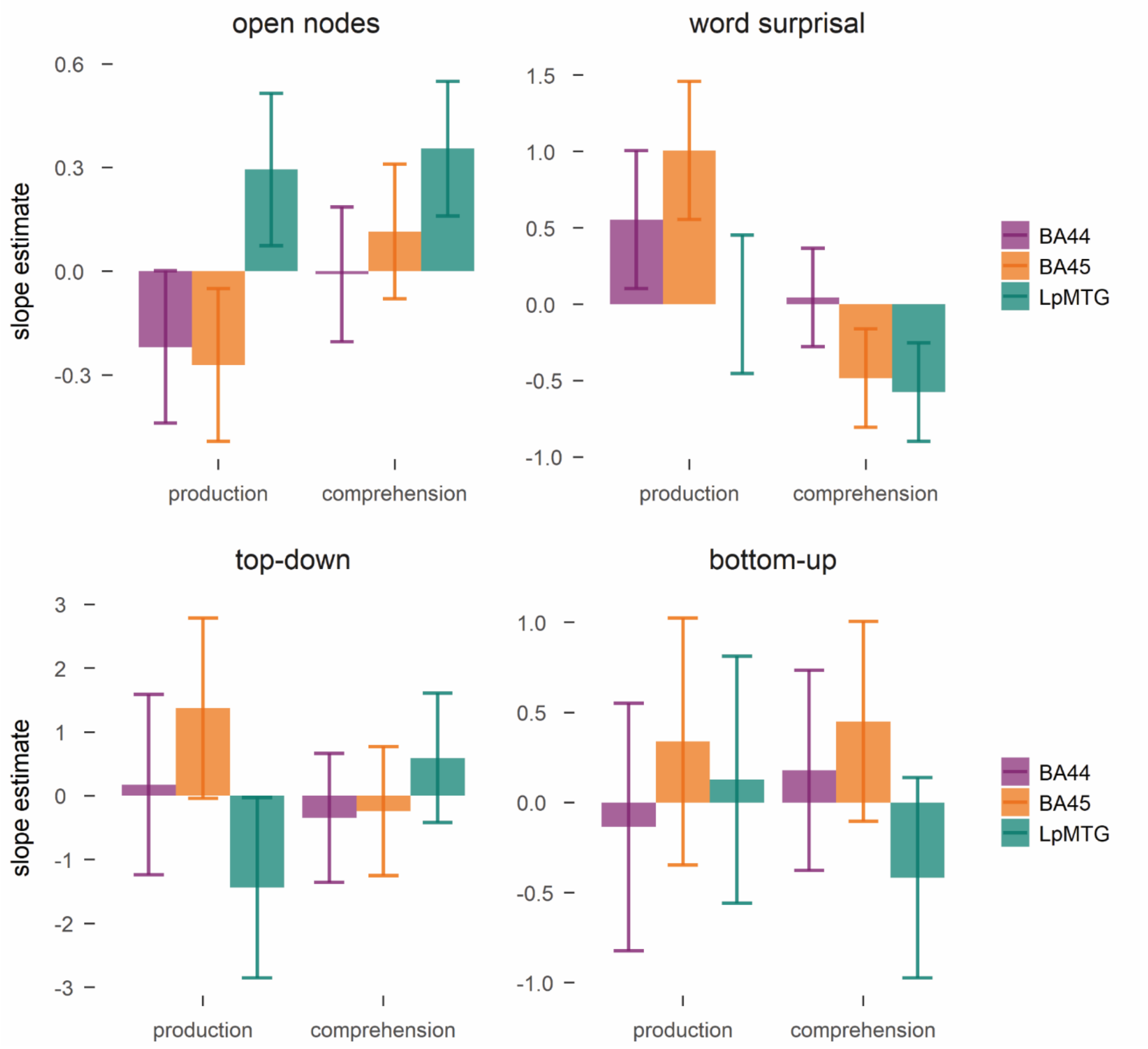


**Supplementary Figure 2**: *Beta estimates for the effect of the temporal derivative of each predictor of on BOLD activity in the regions of interest. Error bars represent confidence intervals. Positive estimates indicate an earlier BOLD response, negative estimates indicate a later BOLD response.*

### Speech fluency: results and discussion

The number of syllables of a word significantly predicted an increase in word duration (unit is seconds, β = 0.08, SE = 0.004, $t = 22.6$, $\chi^2 = 510.02$, $p < 0.0001$), which was expected, but a decrease in pause length before word articulation (β = -0.02, SE = 0.004, $t = 4.9$, $\chi^2 = 23.9$, $p < 0.0001$). The shorter pause before articulation of long words is possibly due to the longer time available to plan for later words during uttering of a long word. Higher word frequency instead predicted shorter word duration (β = -0.017, SE = 0.001, $t = 15.1$, $\chi^2 = 239.8$, $p < 0.0001$), but did not affect pause length (β = 0.0008). Larger word surprisal increased word duration to a small extent (β = 0.004, SE = 0.0004, $t = 9.6$, $\chi^2 = 91.9$, $p < 0.0001$), and it had a larger positive effect on pause length (β = 0.02, SE = 0.002, $t = 10.4$, $\chi^2 = 109.1$, $p < 0.0001$). Less predictable words based on context thus took longer to be initiated and were uttered for a slightly longer time (4 ms), after accounting for their length.

We also determined how predictors of syntactic complexity related to speech fluency. Top-down node counts predicted the largest decrease in word duration (β = -0.045, SE = 0.002, $t = 20.1$, $\chi^2 = 404.1$, $p < 0.0001$), suggesting that when phrases are opened, information can be conveyed faster, possibly to offload working memory. It also predicted the largest increase in pause length before the word in question is uttered (β = 0.09, SE = 0.008, $t = 10.9$, $\chi^2 = 119.7$, $p < 0.0001$) suggesting that grammatical encoding related to a word is performed before word articulation, and that nodes are built in an anticipatory way. Bottom-up parser operations predicted an opposite pattern. Larger bottom-up counts increased word duration (β = 0.012, SE = 0.0009, $t = 12.6$, $\chi^2 = 159.5$, $p < 0.0001$), but decreased pause length (β = 0.021, SE = 0.002, $t = 11.6$, $\chi^2 = 135.7$, $p < 0.0001$). The shorter pauses suggest that at phrase closing the structure is already computed. Finally, open nodes predicted a significant but very small decrease in word duration (β = -0.002, SE = 0.0004, $t = 5.3$, $\chi^2 = 28.5$, $p < 0.0001$), and a larger decrease in pause length (β = -0.024, SE = 0.002, $t = 14.6$, $\chi^2 = 213.9$, $p < 0.0001$), suggesting easier processing the further along in a sentence. In line with the neuroimaging results, this pattern of results suggests that phrase-structure building happens before word articulation and at phrase-opening, with a decrease in pauses the further along in the sentence.

This was the first study to show an increase in neural activity for words associated with higher surprisal, not only in comprehension but also in production. Many studies showed sensitivity of brain activity to surprisal in language comprehension computed with several models (Shain et al., 2020; Willems et al., 2016). The neural results are in line with the behavioural results that show an increase in pause length before less probable words and a small increase in their duration, as found previously (Aylett & Turk, 2004). The results thus converge in demonstrating the sensitivity of the production system to the statistical probabilities of the linguistic input and output, both in behavioural and neural patterns. This finding is in line with accounts of efficient language production that propose a uniform distribution of information in discourse (Uniform Information Density, Jaeger, 2010; Jaeger & Levy, 2007; Karimi, 2022; Piantadosi et al., 2011). More informative units (in information-theoretic terms, i.e. larger surprisal in the current study) take more time in discourse, while redundant units can be uttered faster or eliminated (e.g. for optional words like complementizer *that*, Jaeger, 2010).

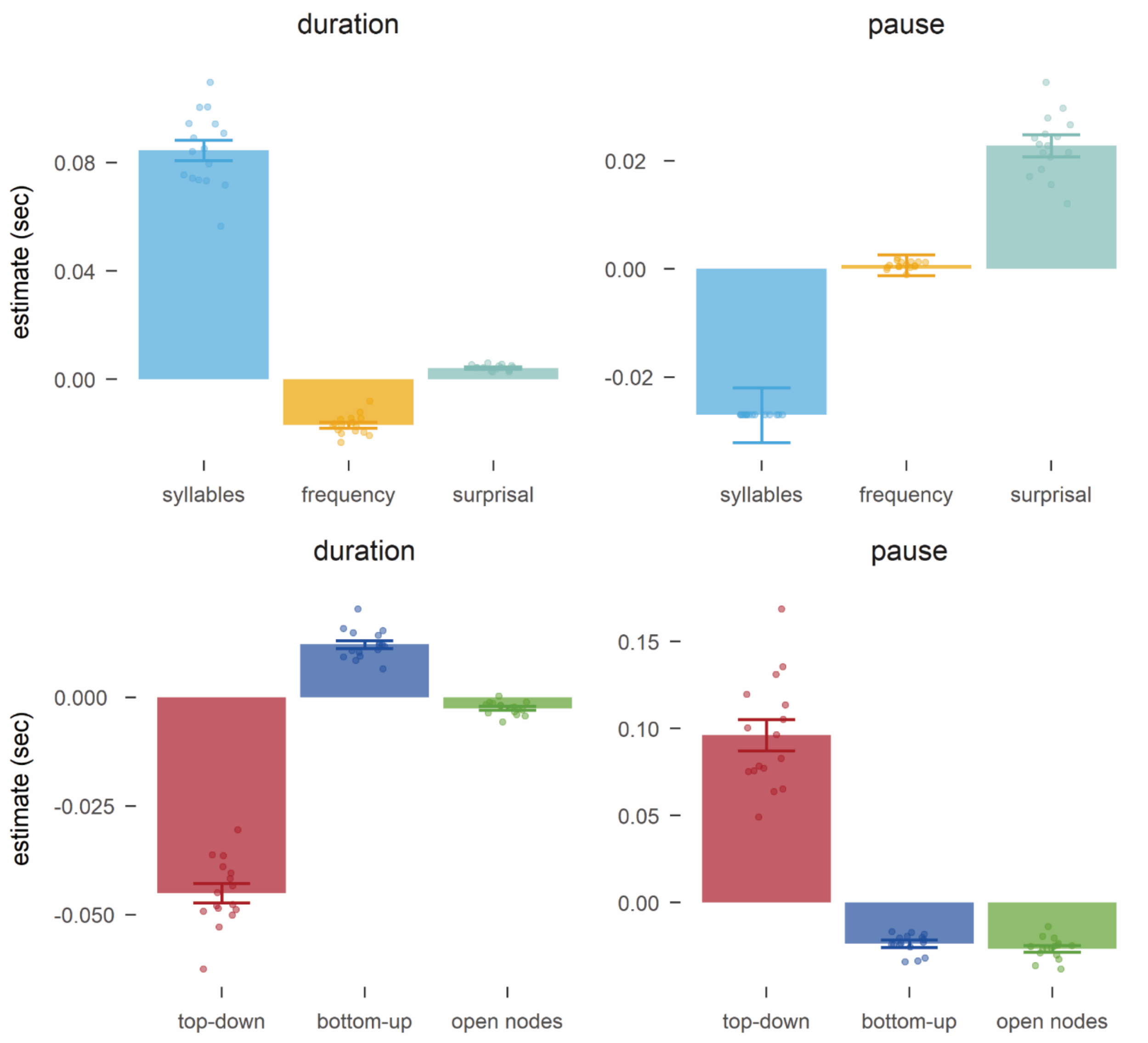


**Supplementary Figure 3:** *Estimates in seconds of the effect of each predictor of word characteristics and phrase-structure building on word durations and pause length before word articulation. Error bars represent standard error of the mean. Individual points represent each participant's estimate as estimated by the random slopes. The model estimated identical random slopes for number of syllables on pause length for each participant.*

### Correlations between model predictors

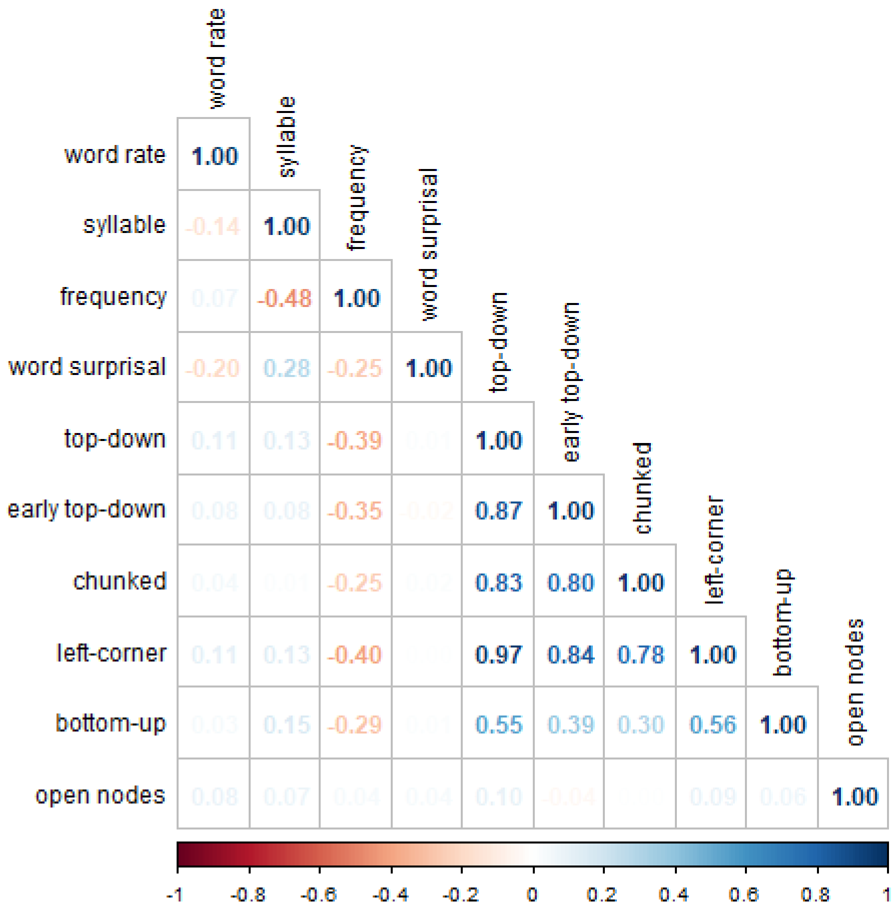


**Supplementary Figure 4**: *Correlation matrix showing Pearson's r correlation among all predictors. Note that not all predictors were used in the same model.*

### Supplementary Table 1

*Summary of model output of BOLD activity in production and comprehension. ROI1 refers to the contrast BA44 vs. BA45, ROI2 refers to the contrast BA44 & BA45 vs. pMTG. Mod stands for modality. AIC stands for Akaike Information Criterion, used for the production only models to determine model fit.*

| | | | **BOLD** | | |
|---|---|---|---|---|---|
| *Predictors* | *Estimates* | *std. Error* | *CI* | *Statistic* | *p* |
| (Intercept) | -0.18 | 0.01 | -0.21 – -0.16 | -14.93 | **<0.001** |
| word rate | 0.54 | 0.03 | 0.48 – 0.61 | 16.15 | **<0.001** |
| syllable | 0.22 | 0.12 | -0.02 – 0.45 | 1.82 | 0.069 |
| frequency | 0.12 | 0.05 | 0.03 – 0.21 | 2.50 | **0.012** |
| top-down | 0.02 | 0.16 | -0.29 – 0.33 | 0.13 | 0.899 |
| ROI1 | 0.00 | 0.01 | -0.01 – 0.01 | 0.26 | 0.796 |
| ROI2 | -0.00 | 0.00 | -0.01 – 0.01 | -0.02 | 0.980 |
| mod1 | -0.01 | 0.00 | -0.02 – 0.00 | -1.81 | 0.070 |
| bottom-up | -0.05 | 0.08 | -0.21 – 0.11 | -0.60 | 0.547 |
| surprisal | 0.16 | 0.02 | 0.12 – 0.21 | 6.67 | **<0.001** |
| open nodes | 0.03 | 0.01 | 0.00 – 0.06 | 2.06 | **0.040** |
| top-down * ROI1 | -0.09 | 0.09 | -0.27 – 0.08 | -1.04 | 0.299 |
| top-down * ROI2 | -0.23 | 0.05 | -0.33 – -0.13 | -4.42 | **<0.001** |
| top-down * mod1 | -0.49 | 0.16 | -0.79 – -0.18 | -3.13 | **0.002** |
| ROI1 * mod1 | 0.00 | 0.01 | -0.01 – 0.01 | 0.06 | 0.956 |
| ROI2 * mod1 | -0.00 | 0.00 | -0.01 – 0.01 | -0.11 | 0.909 |
| ROI1 * bottom-up | 0.05 | 0.06 | -0.07 – 0.17 | 0.81 | 0.420 |
| ROI2 * bottom-up | 0.12 | 0.03 | 0.05 – 0.19 | 3.38 | **0.001** |
| mod1 * bottom-up | 0.39 | 0.08 | 0.23 – 0.55 | 4.84 | **<0.001** |

| | | | | | |
|---|---|---|---|---|---|
| mod1 * surprisal | 0.00 | 0.02 | -0.04 – 0.05 | 0.17 | 0.868 |
| ROI1 * surprisal | 0.08 | 0.02 | 0.04 – 0.12 | 3.63 | **<0.001** |
| ROI2 * surprisal | 0.01 | 0.01 | -0.01 – 0.04 | 0.92 | 0.359 |
| ROI1 * open nodes | 0.02 | 0.01 | 0.00 – 0.03 | 2.02 | **0.044** |
| ROI2 * open nodes | 0.01 | 0.00 | -0.00 – 0.02 | 1.69 | 0.091 |
| mod1 * open nodes | 0.03 | 0.01 | 0.00 – 0.06 | 2.15 | **0.032** |
| top-down * ROI1 * mod1 | -0.20 | 0.09 | -0.37 – -0.02 | -2.21 | **0.027** |
| top-down * ROI2 * mod1 | -0.12 | 0.05 | -0.22 – -0.02 | -2.36 | **0.019** |
| (ROI1 * mod1) * bottom-up | 0.15 | 0.06 | 0.03 – 0.27 | 2.47 | **0.013** |
| (ROI2 * mod1) * bottom-up | 0.09 | 0.03 | 0.02 – 0.16 | 2.56 | **0.010** |
| (ROI1 * mod1) * surprisal | 0.01 | 0.02 | -0.03 – 0.05 | 0.61 | 0.540 |
| (ROI2 * mod1) * surprisal | -0.02 | 0.01 | -0.04 – 0.00 | -1.65 | 0.100 |
| (ROI1 * mod1) * open nodes | 0.02 | 0.01 | 0.00 – 0.03 | 2.54 | **0.011** |
| (ROI2 * mod1) * open nodes | 0.01 | 0.00 | -0.00 – 0.02 | 1.58 | 0.114 |
| $N_{subj}$ | 52 | | | | |
| Observations | 115743 | | | | |
| AIC | 418491.644 | | | | |

### Supplementary Table 2

*Summary of model output of BOLD activity in production with top-down predictor. ROI1 refers to the contrast BA44 vs. BA45, ROI2 refers to the contrast BA44 & BA45 vs. pMTG. Mod stands for modality. AIC stands for Akaike Information Criterion, used for the production only models to determine model fit.*

| | BOLD | | | | |
|---|---|---|---|---|---|
| *Predictors* | *Estimates* | *std. Error* | *CI* | *Statistic* | *p* |
| (Intercept) | -0.15 | 0.02 | -0.18 – -0.11 | -7.56 | **<0.001** |
| word rate | 0.46 | 0.05 | 0.35 – 0.56 | 8.33 | **<0.001** |
| syllable | 0.07 | 0.14 | -0.21 – 0.35 | 0.50 | 0.620 |
| frequency | 0.08 | 0.07 | -0.06 – 0.22 | 1.15 | 0.250 |
| surprisal | 0.15 | 0.03 | 0.09 – 0.22 | 4.78 | **<0.001** |
| ROI1 | 0.00 | 0.01 | -0.02 – 0.02 | 0.12 | 0.905 |
| ROI2 | 0.00 | 0.01 | -0.01 – 0.01 | 0.05 | 0.958 |
| top-down | 0.51 | 0.19 | 0.14 – 0.88 | 2.73 | **0.006** |
| bottom-up | -0.42 | 0.10 | -0.61 – -0.23 | -4.41 | **<0.001** |
| surprisal * ROI1 | 0.06 | 0.04 | -0.01 – 0.14 | 1.75 | 0.080 |
| surprisal * ROI2 | 0.03 | 0.02 | -0.01 – 0.07 | 1.49 | 0.136 |
| ROI1 * top-down | 0.10 | 0.15 | -0.20 – 0.40 | 0.67 | 0.501 |
| ROI2 * top-down | -0.11 | 0.09 | -0.28 – 0.07 | -1.22 | 0.224 |
| ROI1 * bottom-up | -0.10 | 0.10 | -0.29 – 0.09 | -1.03 | 0.303 |
| ROI2 * bottom-up | 0.03 | 0.06 | -0.08 – 0.14 | 0.50 | 0.617 |
| $N_{subj}$ | 16 | | | | |
| Observations | 45099 | | | | |
| AIC | 170821.271 | | | | |

### Supplementary Table 3

*Summary of model output of BOLD activity in production with early top-down predictor. ROI1 refers to the contrast BA44 vs. BA45, ROI2 refers to the contrast BA44 & BA45 vs. pMTG. Mod stands for modality. AIC stands for Akaike Information Criterion, used for the production only models to determine model fit.*

| | BOLD | | | | |
|---|---|---|---|---|---|
| *Predictors* | *Estimates* | *std. Error* | *CI* | *Statistic* | *p* |
| (Intercept) | -0.15 | 0.02 | -0.19 – -0.11 | -7.88 | **<0.001** |
| word rate | 0.47 | 0.05 | 0.37 – 0.58 | 8.69 | **<0.001** |
| syllable | 0.07 | 0.14 | -0.21 – 0.35 | 0.49 | 0.622 |
| frequency | 0.06 | 0.07 | -0.08 – 0.21 | 0.90 | 0.370 |
| surprisal | 0.16 | 0.03 | 0.09 – 0.22 | 4.83 | **<0.001** |
| ROI1 | 0.00 | 0.01 | -0.02 – 0.02 | 0.12 | 0.906 |
| ROI2 | 0.00 | 0.01 | -0.01 – 0.01 | 0.06 | 0.956 |
| early top-down | 0.33 | 0.20 | -0.06 – 0.72 | 1.64 | 0.100 |
| bottom-up | -0.35 | 0.10 | -0.55 – -0.16 | -3.54 | **<0.001** |
| surprisal * ROI1 | 0.06 | 0.04 | -0.01 – 0.14 | 1.78 | 0.076 |
| surprisal * ROI2 | 0.03 | 0.02 | -0.01 – 0.07 | 1.42 | 0.155 |
| ROI1 * early top-down | 0.10 | 0.12 | -0.14 – 0.35 | 0.85 | 0.397 |
| ROI2 * early top-down | -0.17 | 0.07 | -0.31 – -0.03 | -2.34 | **0.020** |
| ROI1 * bottom-up | -0.09 | 0.09 | -0.27 – 0.08 | -1.04 | 0.296 |
| ROI2 * bottom-up | 0.04 | 0.05 | -0.07 – 0.14 | 0.68 | 0.494 |
| $N_{subj}$ | 16 | | | | |
| Observations | 45099 | | | | |
| AIC | 170803.949 | | | | |

### Supplementary Table 4

*Summary of model output of BOLD activity in production with chunked top-down predictor. ROI1 refers to the contrast BA44 vs. BA45, ROI2 refers to the contrast BA44 & BA45 vs. pMTG. Mod stands for modality. AIC stands for Akaike Information Criterion, used for the production only models to determine model fit.*

| | **BOLD** | | | | |
|---|---|---|---|---|---|
| *Predictors* | *Estimates* | *std. Error* | *CI* | *Statistic* | *p* |
| (Intercept) | -0.16 | 0.02 | -0.19 – -0.12 | -8.13 | **<0.001** |
| word rate | 0.49 | 0.05 | 0.38 – 0.59 | 8.97 | **<0.001** |
| syllable | 0.07 | 0.14 | -0.21 – 0.35 | 0.52 | 0.603 |
| frequency | 0.05 | 0.07 | -0.09 – 0.19 | 0.69 | 0.491 |
| surprisal | 0.15 | 0.03 | 0.09 – 0.21 | 4.62 | **<0.001** |
| ROI1 | 0.00 | 0.01 | -0.02 – 0.02 | 0.12 | 0.906 |
| ROI2 | 0.00 | 0.01 | -0.01 – 0.01 | 0.05 | 0.957 |
| chunked top-down | 0.32 | 0.16 | 0.00 – 0.63 | 1.97 | **0.049** |
| bottom-up | -0.31 | 0.08 | -0.47 – -0.14 | -3.65 | **<0.001** |
| surprisal * ROI1 | 0.06 | 0.04 | -0.01 – 0.14 | 1.75 | 0.080 |
| surprisal * ROI2 | 0.03 | 0.02 | -0.01 – 0.07 | 1.49 | 0.135 |
| ROI1 * chunked top-down | 0.02 | 0.13 | -0.23 – 0.27 | 0.17 | 0.866 |
| ROI2 * chunked top-down | -0.04 | 0.07 | -0.19 – 0.10 | -0.56 | 0.574 |
| ROI1 * bottom-up | -0.07 | 0.09 | -0.25 – 0.10 | -0.82 | 0.413 |
| ROI2 * bottom-up | 0.00 | 0.05 | -0.10 – 0.10 | 0.02 | 0.984 |
| $N_{subj}$ | 16 | | | | |
| Observations | 45099 | | | | |
| AIC | 170834.489 | | | | |

### Supplementary Table 5

*Summary of model output of BOLD activity in production and comprehension, including the temporal derivative (der) of all predictors of interest. ROI1 refers to the contrast BA44 vs. BA45, ROI2 refers to the contrast BA44 & BA45 vs. pMTG. Mod stands for modality. AIC stands for Akaike Information Criterion, used for the production only models to determine model fit.*

| | BOLD | | | | |
|---|---|---|---|---|---|
| *Predictors* | *Estimates* | *std. Error* | *CI* | *Statistic* | *p* |
| (Intercept) | -0.17 | 0.01 | -0.20 – -0.15 | -13.99 | **<0.001** |
| word rate | 0.52 | 0.03 | 0.45 – 0.58 | 15.12 | **<0.001** |
| syll | 0.22 | 0.12 | -0.00 – 0.45 | 1.93 | 0.054 |
| frequency | 0.12 | 0.05 | 0.03 – 0.21 | 2.52 | **0.012** |
| top-down der | 0.02 | 0.35 | -0.66 – 0.71 | 0.06 | 0.953 |
| ROI1 | 0.00 | 0.01 | -0.01 – 0.01 | 0.26 | 0.795 |
| ROI2 | 0.00 | 0.00 | -0.01 – 0.01 | 0.00 | 0.999 |
| mod1 | -0.01 | 0.00 | -0.02 – 0.00 | -1.72 | 0.085 |
| bottom-up der | 0.09 | 0.16 | -0.23 – 0.41 | 0.56 | 0.576 |
| surprisal der | 0.09 | 0.12 | -0.14 – 0.33 | 0.76 | 0.448 |
| open nodes der | 0.04 | 0.05 | -0.06 – 0.15 | 0.84 | 0.399 |
| top-down | -0.10 | 0.16 | -0.42 – 0.21 | -0.62 | 0.533 |
| bottom-up | 0.07 | 0.10 | -0.13 – 0.27 | 0.65 | 0.515 |
| surprisal | 0.16 | 0.02 | 0.11 – 0.21 | 6.54 | **<0.001** |
| open nodes | 0.03 | 0.02 | -0.00 – 0.06 | 1.81 | 0.071 |
| top-down der * ROI1 | 0.33 | 0.24 | -0.14 – 0.79 | 1.38 | 0.166 |
| top-down der * ROI2 | -0.22 | 0.14 | -0.49 – 0.05 | -1.62 | 0.105 |
| top-down der * mod1 | -0.02 | 0.35 | -0.70 – 0.67 | -0.05 | 0.962 |

| | | | | | |
|---|---|---|---|---|---|
| ROI1 * mod1 | 0.00 | 0.01 | -0.01 – 0.01 | 0.06 | 0.952 |
| ROI2 * mod1 | -0.00 | 0.00 | -0.01 – 0.01 | -0.12 | 0.906 |
| ROI1 * bottom-up der | 0.19 | 0.14 | -0.08 – 0.45 | 1.37 | 0.171 |
| ROI2 * bottom-up der | -0.12 | 0.08 | -0.27 – 0.04 | -1.50 | 0.134 |
| mod1 * bottom-up der | -0.02 | 0.16 | -0.33 – 0.30 | -0.12 | 0.904 |
| mod1 * surprisal der | -0.43 | 0.12 | -0.66 – -0.19 | -3.56 | **<0.001** |
| ROI1 * surprisal der | -0.02 | 0.06 | -0.15 – 0.11 | -0.29 | 0.775 |
| ROI2 * surprisal der | -0.19 | 0.04 | -0.26 – -0.12 | -5.08 | **<0.001** |
| ROI1 * open nodes der | 0.02 | 0.05 | -0.07 – 0.11 | 0.39 | 0.700 |
| ROI2 * open nodes der | 0.14 | 0.03 | 0.09 – 0.19 | 5.22 | **<0.001** |
| mod1 * open nodes der | 0.11 | 0.05 | 0.01 – 0.21 | 2.09 | **0.037** |
| ROI1 * top-down | -0.13 | 0.13 | -0.38 – 0.12 | -1.05 | 0.295 |
| ROI2 * top-down | -0.49 | 0.07 | -0.63 – -0.34 | -6.63 | **<0.001** |
| mod1 * top-down | -0.67 | 0.16 | -0.98 – -0.35 | -4.16 | **<0.001** |
| ROI1 * bottom-up | 0.09 | 0.11 | -0.13 – 0.30 | 0.80 | 0.427 |
| ROI2 * bottom-up | 0.38 | 0.06 | 0.26 – 0.50 | 6.08 | **<0.001** |
| mod1 * bottom-up | 0.57 | 0.10 | 0.37 – 0.77 | 5.54 | **<0.001** |
| mod1 * surprisal | 0.01 | 0.02 | -0.04 – 0.05 | 0.25 | 0.803 |
| ROI1 * surprisal | 0.08 | 0.02 | 0.04 – 0.12 | 3.68 | **<0.001** |
| ROI2 * surprisal | 0.01 | 0.01 | -0.01 – 0.03 | 0.81 | 0.417 |
| ROI1 * open nodes | 0.01 | 0.01 | -0.00 – 0.03 | 1.38 | 0.166 |
| ROI2 * open nodes | 0.01 | 0.00 | 0.00 – 0.02 | 1.98 | **0.048** |
| mod1 * open nodes | 0.03 | 0.02 | 0.00 – 0.06 | 2.04 | **0.041** |
| top-down d * ROI1 * mod1 | -0.27 | 0.24 | -0.74 – 0.19 | -1.16 | 0.248 |
| top-down d * ROI2 * mod1 | 0.52 | 0.14 | 0.25 – 0.78 | 3.79 | **<0.001** |
| (ROI1 * mod1) * bottom-up der | -0.05 | 0.14 | -0.32 – 0.22 | -0.37 | 0.711 |

| | | | | | |
|---|---|---|---|---|---|
| (ROI2 * mod1) * bottom-up der | -0.13 | 0.08 | -0.28 – 0.03 | -1.60 | 0.109 |
| (ROI1 * mod1) * surprisal der | -0.24 | 0.06 | -0.37 – -0.12 | -3.79 | **<0.001** |
| (ROI2 * mod1) * surprisal der | 0.07 | 0.04 | -0.00 – 0.14 | 1.90 | 0.057 |
| (ROI1 * mod1) * open nodes der | 0.04 | 0.05 | -0.05 – 0.14 | 0.94 | 0.345 |
| (ROI2 * mod1) * open nodes der | -0.04 | 0.03 | -0.09 – 0.01 | -1.48 | 0.140 |
| (ROI1 * mod1) * top-down | -0.27 | 0.13 | -0.52 – -0.02 | -2.13 | **0.033** |
| (ROI2 * mod1) * top-down | -0.07 | 0.07 | -0.22 – 0.07 | -1.00 | 0.318 |
| (ROI1 * mod1) * bottom-up | 0.22 | 0.11 | 0.01 – 0.43 | 2.07 | **0.039** |
| (ROI2 * mod1) * bottom-up | 0.02 | 0.06 | -0.10 – 0.15 | 0.39 | 0.696 |
| (ROI1 * mod1) * surprisal | 0.01 | 0.02 | -0.03 – 0.05 | 0.58 | 0.565 |
| (ROI2 * mod1) * surprisal | -0.02 | 0.01 | -0.04 – 0.01 | -1.50 | 0.133 |
| (ROI1 * mod1) * open nodes | 0.02 | 0.01 | 0.00 – 0.04 | 2.30 | **0.021** |
| (ROI2 * mod1) * open nodes | 0.01 | 0.00 | 0.00 – 0.02 | 2.69 | **0.007** |
| $N_{subj}$ | 52 | | | | |
| Observations | 115743 | | | | |
| AIC | 418263.250 | | | | |

### Supplementary Table 6

*Summary of model output of the pause length preceding each word's production. AIC stands for Akaike Information Criterion, used for the production only models to determine model fit.*

| | pause length | | | | |
|---|---|---|---|---|---|
| *Predictors* | *Estimates* | *std. Error* | *CI* | *Statistic* | *p* |
| (Intercept) | 0.15 | 0.02 | 0.11 – 0.19 | 7.11 | **<0.001** |
| frequency | 0.00 | 0.00 | -0.00 – 0.00 | 0.36 | 0.722 |
| surprisal | 0.02 | 0.00 | 0.02 – 0.03 | 11.22 | **<0.001** |
| syllables | -0.03 | 0.01 | -0.04 – -0.02 | -5.36 | **<0.001** |
| bottom-up | -0.02 | 0.00 | -0.03 – -0.02 | -10.36 | **<0.001** |
| open nodes | -0.03 | 0.00 | -0.03 – -0.02 | -13.66 | **<0.001** |
| top-down | 0.10 | 0.01 | 0.08 – 0.11 | 10.66 | **<0.001** |
| $N_{subj}$ | 16 | | | | |
| Observations | 45079 | | | | |
| AIC | 82830.780 | | | | |

### Supplementary Table 7

*Summary of model output of word duration. AIC stands for Akaike Information Criterion, used for the production only models to determine model fit.*

| | word duration | | | | |
|---|---|---|---|---|---|
| *Predictors* | *Estimates* | *std. Error* | *CI* | *Statistic* | *p* |
| (Intercept) | 0.27 | 0.01 | 0.25 – 0.29 | 28.22 | **<0.001** |
| frequency | -0.02 | 0.00 | -0.02 – -0.01 | -15.44 | **<0.001** |
| surprisal | 0.00 | 0.00 | 0.00 – 0.01 | 9.38 | **<0.001** |
| syllables | 0.08 | 0.00 | 0.08 – 0.09 | 22.50 | **<0.001** |
| bottom-up | 0.01 | 0.00 | 0.01 – 0.01 | 12.61 | **<0.001** |
| open nodes | -0.00 | 0.00 | -0.00 – -0.00 | -5.35 | **<0.001** |
| top-down | -0.05 | 0.00 | -0.05 – -0.04 | -20.05 | **<0.001** |
| $N_{subj}$ | 16 | | | | |
| Observations | 45079 | | | | |
| AIC | -42830.283 | | | | |

### References


Aylett, M., & Turk, A. (2004). The Smooth Signal Redundancy Hypothesis: A Functional Explanation for Relationships between Redundancy, Prosodic Prominence, and Duration in Spontaneous Speech. *Language and Speech*, *47*(1), 31–56. https://doi.org/10.1177/00238309040470010201

Jaeger, T. F. (2010). Redundancy and reduction: Speakers manage syntactic information density. *Cognitive Psychology*, *61*(1), 23–62. https://doi.org/10.1016/j.cogpsych.2010.02.002

Jaeger, T. F., & Levy, R. (2007). Speakers optimize information density through syntactic reduction. *Advances in Neural Information Processing Systems*, *19*. https://proceedings.neurips.cc/paper/2006/hash/c6a01432c8138d46ba39957a8250e027-Abstract.html

Karimi, H. (2022). Greater entropy leads to more explicit referential forms during language production. *Cognition*, *225*, 105093. https://doi.org/10.1016/j.cognition.2022.105093

Piantadosi, S. T., Tily, H., & Gibson, E. (2011). Word lengths are optimized for efficient communication. *Proceedings of the National Academy of Sciences*, *108*(9), 3526–3529. https://doi.org/10.1073/pnas.1012551108

Shain, C., Blank, I. A., van Schijndel, M., Schuler, W., & Fedorenko, E. (2020). FMRI reveals language-specific predictive coding during naturalistic sentence comprehension. *Neuropsychologia*, *138*, 107307. https://doi.org/10.1016/j.neuropsychologia.2019.107307

Willems, R. M., Frank, S. L., Nijhof, A. D., Hagoort, P., & van den Bosch, A. (2016). Prediction During Natural Language Comprehension. *Cerebral Cortex*, *26*(6), 2506–2516. https://doi.org/10.1093/cercor/bhv075
